# Quantification of cellular contributions to multi-cellular pattern formation^*^

**DOI:** 10.64898/2026.10.08.757240

**Authors:** Maxime Estavoyer, Michael L Zhao, Matthias Merkel

## Abstract

Multi-cellular pattern formation is a key process in organism development, regeneration, and disease. Such multi-cellular patterns include, for instance, cell density and gene expression patterns. Here, we develop a cell-track-based method that quantifies how much individual cellular processes contribute to the emergence of large-scale patterns. These cellular processes include changes in cellular expression levels, cell motion, cell divisions, and apoptosis. We validate our method using synthetic data, before demonstrating it on two sets of experimental data. Our method is purely geometric and does not require any mechanistic assumptions about the underlying pattern-forming processes, making it very generic and robust. It thus enables unbiased, data-driven study of the mechanisms driving pattern formation across a broad class of systems.

## I. INTRODUCTION

Multi-cellular pattern formation is a key process during development, regeneration, and disease [1–4]. Recent examples of pattern formation during animal development include patterning of skin follicles [5–8], vertebrate digit patterning [9, 10], pigment pattern formation [11–14], tooth patterning [15, 16], and the patterning of stem cell aggregates [17–20]. Patterning during regeneration is observed for instance in planarians, salamanders, and hydra [21–23]. Finally, multi-cellular patterns also appear, for instance, in skin diseases [4] and cancer [4, 24]. Yet, dissecting the mechanisms driving patterning remains challenging.

Pattern formation can be driven by different cellular processes, which include cell differentiation, cell motility, divisions, and apoptosis. Sometimes only one of these processes dominates. For instance, in many systems pattern formation is believed to be driven predominantly by cell differentiation dynamics, often mediated by morphogens [9, 10, 18, 25–27]. In other cases, the main driver of pattern formation is cell motility [5, 7, 28]. Yet, it becomes clearer that pattern formation is often driven by a combination of different cellular processes [19, 29–31]. Furthermore, sometimes the emergent pattern is always of the same type (e.g. stripes, a single dot, several dots) [32–34], while in other cases the type of emergent pattern depends on system size [17, 20]. So far, we are missing a generally applicable way of quantifying pattern formation and the cellular contributions to it, independently of the precise type of pattern.

To pinpoint the mechanism driving pattern formation, most often researchers follow a *hypothesis-driven* approach [5–7, 9–11, 13, 15, 18, 30, 35–38], where models are postulated that make specific assumptions about which cellular processes drive pattern formation and how these processes function. Predictions of these models are then compared to experimental data [39].

Alternatively, there exist a few *data-driven* approaches to quantify multi-cellular patterns directly from imaging data, including most notably pair-correlation functions and related spatial statistics [40–44]. The advantage of data-driven approaches is that they usually do not require any prior knowledge or assumptions about the system under study. However, so far, these approaches do not disentangle how patterns emerge as a result of different cellular processes.

Here, we propose a data-driven framework that fills this gap, leveraging recent improvements in live imaging and automated cell tracking [45–47]. Based on cell tracks and gene expression information, we quantify measures for multi-cellular pattern formation, and decompose them into contributions by cellular processes. This quantification is automated by our python tool CREDIT (Cellular Roles in Emergent Development Inferred from Tracks). We validate our approach using agent-based models that allow us to tune how much the different cellular processes contribute to pattern formation. We then apply our framework to experimental data. First, in quasi-2D aggregates of mouse presomitic mesoderm cells, patterns of the mesodermal marker T/Brachyury emerge [20], and our analysis reveals that this is primarily driven by cell motility rather than gene expression. Second, breast cancer cell populations display the emergence of a pattern of cancer stem cell domains, akin to niches [24]. Our method shows that the formation of these domains is mostly driven by cell divisions with a negative net contribution by gene expression.

Our approach is purely geometric, i.e. it does not require any mechanistic assumption about the underlying pattern forming process. It can be applied to any set of experimental or simulation data that provides cell tracks and expression data. Finally, because our method allows to pinpoint the cellular processes driving pattern formation, it can provide the basis for any subsequent hypothesis-driven refinement.

## II. RESULTS

### A. Measures for multi-cellular pattern formation

#### 1. Definitions

We start from a cell population of time-dependent size *N* (*t*). The cells, labeled by *i* = 1, …, *N* (*t*), have known time-dependent 2D positions ***r***_*i*_(*t*) and expression levels *f*_*i*_(*t*) of some protein. Both can be quantified from live imaging data; positions are provided by tracking software, while expression levels can be quantified as the fluorescence intensity integrated over the cell or nucleus volume. Note that more generally, the *f*_*i*_(*t*) data can represent any cell-associated quantity, such as for instance the cell area or a superposition of several protein expression levels.

Using the ***r***_*i*_(*t*) and *f*_*i*_(*t*) data, we compute measures for different types of multi-cellular patterns (Figure 1A). First, we consider the cell number density pattern *ρ*(***r***, *t*) (Figure 1Ai). The fact that it emerges from individual cell positions is formally written as 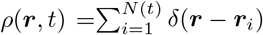 where *δ* denotes the Dirac distribution. Second, one could consider some cell expression pattern, i.e. some continuous spatial function *f* (***r***, *t*) that corresponds to the measured cellular expression levels at each cell position (Figure 1Aii), i.e. *f* (***r***_*i*_, *t*) = *f*_*i*_. Yet, strictly speaking, the value of *f* (***r***, *t*) away from any cell position is unknown. Finally, microscopy images most directly show an apparent expression pattern – i.e. protein expression exists only where cells are (Figure 1Aiii). This pattern corresponds to the product 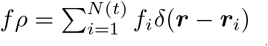 Here, we focus on quantifying (i) the cell number density pattern *ρ* and (ii) the apparent expression pattern *fρ*.

**Figure 1.**
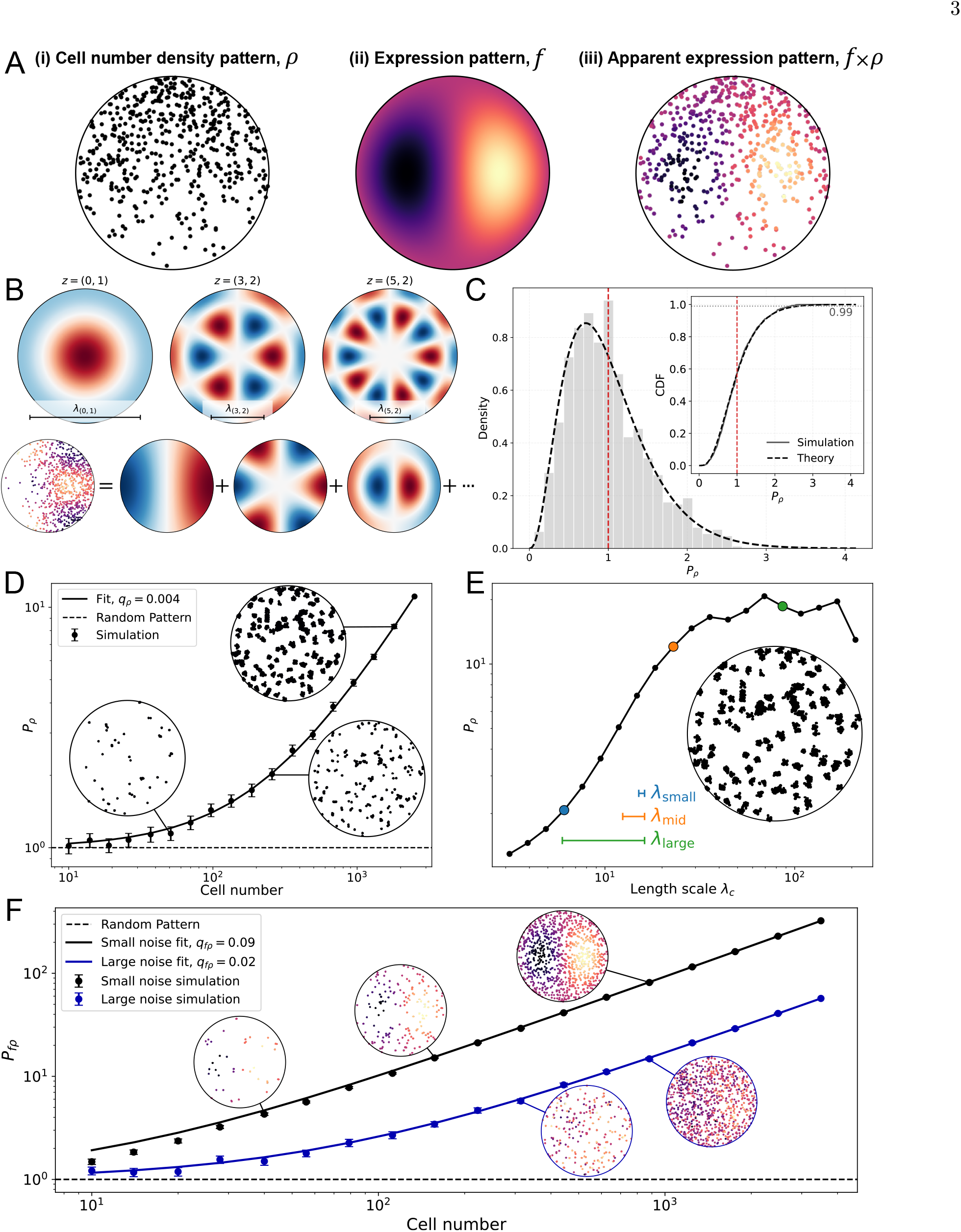
Quantification of multi-cellular patterns from cell positions and expression data. (A) Different multi-cellular patterns can be considered: (i) cell number density pattern *ρ*(***r***, *t*), (ii) continuous expression pattern *f* (***r***, *t*), and (iii) apparent expression pattern *fρ*. (B) Example Fourier–Bessel modes *ϕ*_*z*_ (***r***) associated with characteristic length scales *λ*_*z*_, and decomposition of an apparent expression pattern into a weighted sum of these modes. (C) Distribution of the density pattern measure *P*_*ρ*_ for randomly distributed cells. Gray bars indicate simulated data and the black dashed curve is an analytical prediction, whose average is marked by the red dashed line. Here, *M*_*c*_ = 7 modes were taken into account to compute *P*_*ρ*_. (inset) Same data plotted as cumulative distribution, from which significance levels can be directly read off. (D) Scaling of the density pattern measure *P*_*ρ*_ with cell number *N*. Bootstrapped subsets of cells follow the predicted linear scaling in Eq. (5). (E) Dependence of the density pattern measure *P*_*ρ*_ on the cutoff length scale *λ*_*c*_. (F) Scaling of the apparent expression pattern measure *P*_*fρ*_ with cell number *N* for spatial expression patterns with low and high expression noise. Bootstrapped subsets of cells follow the predicted linear scaling in Eq. (6).

To compute a measure for pattern formation in the fields *ρ* or *fρ*, we quantify their *variability on large length scales*. To this end, we first express these fields in some orthonormal basis (Figure 1B). For all cases studied here, we use a so-called Fourier-Bessel transformation, which acts on a circular domain of radius *R* (see SI section 2). The advantage of such a transformation is that the overall variability (e.g. the variance in the apparent spatial expression pattern) can be expressed in a way that depends on the length scale *λ*. More precisely, the fields *ρ* or *fρ* are decomposed into a sum of basis functions *ϕ*_*z*_(***r***), each of which varies on a different length scale *λ*_*z*_ (Figure 1B):

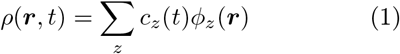

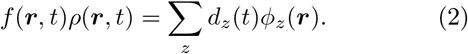

The coefficients *c*_*z*_(*t*) and *d*_*z*_(*t*) express at each time point how much the respective basis function *ϕ*_*z*_(***r***) contributes to the patterns *ρ* and *fρ*. Based on these decompositions, we introduce our measures for pattern formation, *P*_*ρ*_(*t*) and *P*_*fρ*_(*t*), as the *variance created by the modes ϕ*_*z*_(***r***) *with the largest length scales*

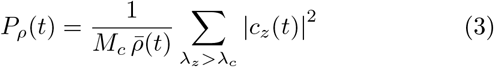

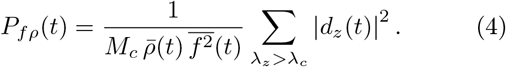

Here, the sums are over the mode index *z*, excluding the spatially homogeneous mode *z* = (0, 0). The lower length scale limit *λ*_*c*_ is a cutoff parameter that defines above what length scale variability is considered not as a local fluctuation but as contributing to the multi-cellular pattern. In Eqs. (3) and (4), we normalize the measures by *M*_*c*_, which denotes the number of non-homogeneous modes with *λ*_*z*_ *> λ*_*c*_, by the average cell number density 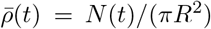, and by the average squared cellular expression level 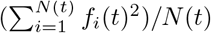.These normalizations ensure that randomly placed cells with random expression levels yield on average *P*_*ρ*_ = *P*_*fρ*_ = 1. However, for any systematic pattern in the cell density *ρ* or the apparent expression *fρ*, the respective measure *P*_*ρ*_ or *P*_*fρ*_ becomes larger than one (see next section and SI section 3).

#### 2. Validation

We use simulated data to test whether both measures, *P*_*ρ*_ and *P*_*fρ*_, reliably indicate the presence or absence of a pattern.

To this end, we first study the values that these measures take when used on random data, where we start by testing the cell number density pattern measure *P*_*ρ*_. Specifically, we randomly distribute *N* cell positions ***r***_*i*_ independently and uniformly on the disk with radius *R*, and then compute the pattern measure *P*_*ρ*_. Note that although the cells are uniformly distributed, the pattern measure *P*_*ρ*_ will be small, but not zero, since fluctuations always occur. We thus simulate the distribution of values *P*_*ρ*_ that one would obtain for many randomly generated patterns, and we find that it matches an analytical prediction derived in the SI (compare histogram to black dashed curve in Figure 1C). As mentioned before, the average of the distribution is *P*_*ρ*_ = 1. Hence, *P*_*ρ*_ attains the expected value of *P*_*ρ*_ ~1 for random data.

Yet, for random data, *P*_*ρ*_ will not be *exactly* one, but fluctuate around it. If one measures some value *P*_*ρ*_ *≳* 1, it would be good to have a statistical test saying how likely it is that we just look at noise. To this end, we plot the cumulative distribution of the value *P*_*ρ*_ in the inset to Figure 1C. For instance, if the cells were just randomly distributed (null hypothesis), this plot tells us that we would measure *P*_*ρ*_ value smaller than 2.6 with a probability of 99%. In other words, for *P*_*ρ*_ *>* 2.6, there is a confidence of 99% that there is a pattern in the cell number density. This significance level only depends on the number of modes *M*_*c*_ that were taken into account to compute *P*_*ρ*_, i.e. it only depends on the cutoff length scale *λ*_*c*_: The smaller *λ*_*c*_, the more modes *M*_*c*_ are included, the narrower the distribution in Figure 1C, and the more restrictive the significance levels (Supplemental Figure S2). Taken together, a convenient feature of our method is that statistical significance is directly provided by the value of our measure itself.

We next test the apparent expression pattern measure *P*_*fρ*_ on random data. We randomly distribute cell positions as before, and additionally randomly pick the expression levels *f*_*i*_ independent of the cell positions. Again, the resulting simulated distribution perfectly matches the predicted distribution with average *P*_*fρ*_ = 1 (see SI Fig. S1). Furthermore, the value of *P*_*fρ*_ again also acts as measure for statistical significance where *P*_*fρ*_ *>* 2.6 indicates a confidence of 99% that there is an apparent expression pattern (see SI Fig. S1 inset). Note that due to the choice of normalization in Eq. (4) this significance test is independent of any scaling of the expression variables *f*_*i*_. Hence, for random data, both measures, *P*_*ρ*_ and *P*_*fρ*_, reliably indicate the absence of a pattern.

We next test whether conversely, our measures also reliably indicate the presence of a pattern. We first note that in the presence of a pattern, the values of *P*_*ρ*_ and *P*_*fρ*_ depend on the number of cells (see SI for details). Briefly, this is because for the modes *z* that describe the actual pattern, the coefficients *c*_*z*_ and *d*_*z*_ scale linearly with the cell number *N*. Meanwhile, for purely random data in the absence of a pattern, only the squares, |*c*_*z*_| ^2^ and |*d*_*z*_| ^2^, scale linearly with *N*. Thus, for random data, due to the normalization in Eqs. (3) and (4) with 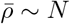, the measures *P*_*ρ*_ and *P*_*fρ*_ do not scale with *N*, and instead have average 1. However, when there is a pattern, even a noisy one, an additional term arises that linearly scales with the cell number *N* :

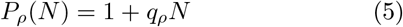

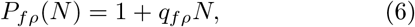

where *q*_*ρ*_ and *q*_*fρ*_ are prefactors, such that *q*_*ρ*_*N* and *q*_*fρ*_*N* can be interpreted as signal-to-noise ratios.

We now test these ideas on noisy patterns. We test the cell density pattern measure *P*_*ρ*_ on the cell set in Figure 1D, with 2500 cells. We furthermore vary the cell number *N* by applying a bootstrapping method (SI section 4), i.e. we select random subsets of the full set of cells, each time computing *P*_*ρ*_ of that subset. We find indeed an excellent fit to the prediction in Eq. (5) with *q*_*ρ*_ = 0.004. We further test the apparent expression pattern measure *P*_*fρ*_ on the cell sets in Figure 1F, right insets. These cell sets include low (upper inset) and high (lower inset) cell expression noise. Using again the bootstrapping method, we find again good fits to Eq. (6) with *q*_*fρ*_ = 0.09 (low expression noise) and *q*_*fρ*_ = 0.02 (high expression noise), respectively (Figure 1F). Taken together, both pattern measures reliably indicate the presence of a pattern. Both increase linearly with the number of cells, where the prefactors *q*_*ρ*_ and *q*_*fρ*_ provide the respective signal-to-noise ratio per cell.

Finally, we note that the pattern measures also depend on the cutoff length scale *λ*_*c*_ above which variability is considered part of the pattern as opposed to local fluctuations. For instance, consider the cell arrangement in Figure 1E inset, which is made up of several randomly placed clusters, each consisting of several cells. On length scales larger than the cluster size, this is a random pattern, and many modes with large *λ*_*z*_ contribute to the pattern. However, modes with *λ*_*z*_ much smaller than the cluster size do not substantially contribute. Thus, as the cutoff length scale *λ*_*c*_ is decreased, more and more modes are taken into account, and since we quantify the *average* variation per mode, we observe a decrease of the pattern measure *P*_*ρ*_ as we decrease *λ*_*c*_. However, at larger *λ*_*c*_, we find a plateau indicating that at large length scales different modes contribute similarly to the overall variability. A similar *λ*_*c*_ dependence is also obtained for *P*_*fρ*_ (see SI Fig. S3).

Taken together, both pattern measures reliably indicate presence and absence of patterns, where the definition of what a pattern depends on the cutoff length scale *λ*_*c*_, which separates small-scale noise from large-scale signal.

### B. Cellular contributions to pattern formation

#### 1. Definitions

The pattern measures *P*_*ρ*_(*t*) and *P*_*fρ*_(*t*) generally vary over time, which is due to several cellular processes, including changes of gene expression levels, cell motion, cell divisions, and cell disappearance. To quantify the contributions of these processes to pattern formation, we note that the pattern measures depend only on cell number *N*, cell positions ***r***_*i*_, and expression levels *f*_*i*_. In other words, any change in *P*_*ρ*_ and *P*_*fρ*_ can only be due to changes in cell number *N*, i.e. divisions and cell disappearance, changes in cell positions ***r***_*i*_, i.e. cell motion, and changes in expression levels *f*_*i*_. Based on these considerations, we can uniquely decompose changes in *P*_*ρ*_ and *P*_*fρ*_ into cellular contributions (Figure 2A, Figure 3A, details in SI section 5). Specifically, between two subsequent time points, *t*_*n*_ *→ t*_*n*+1_, we write:

**Figure 2.**
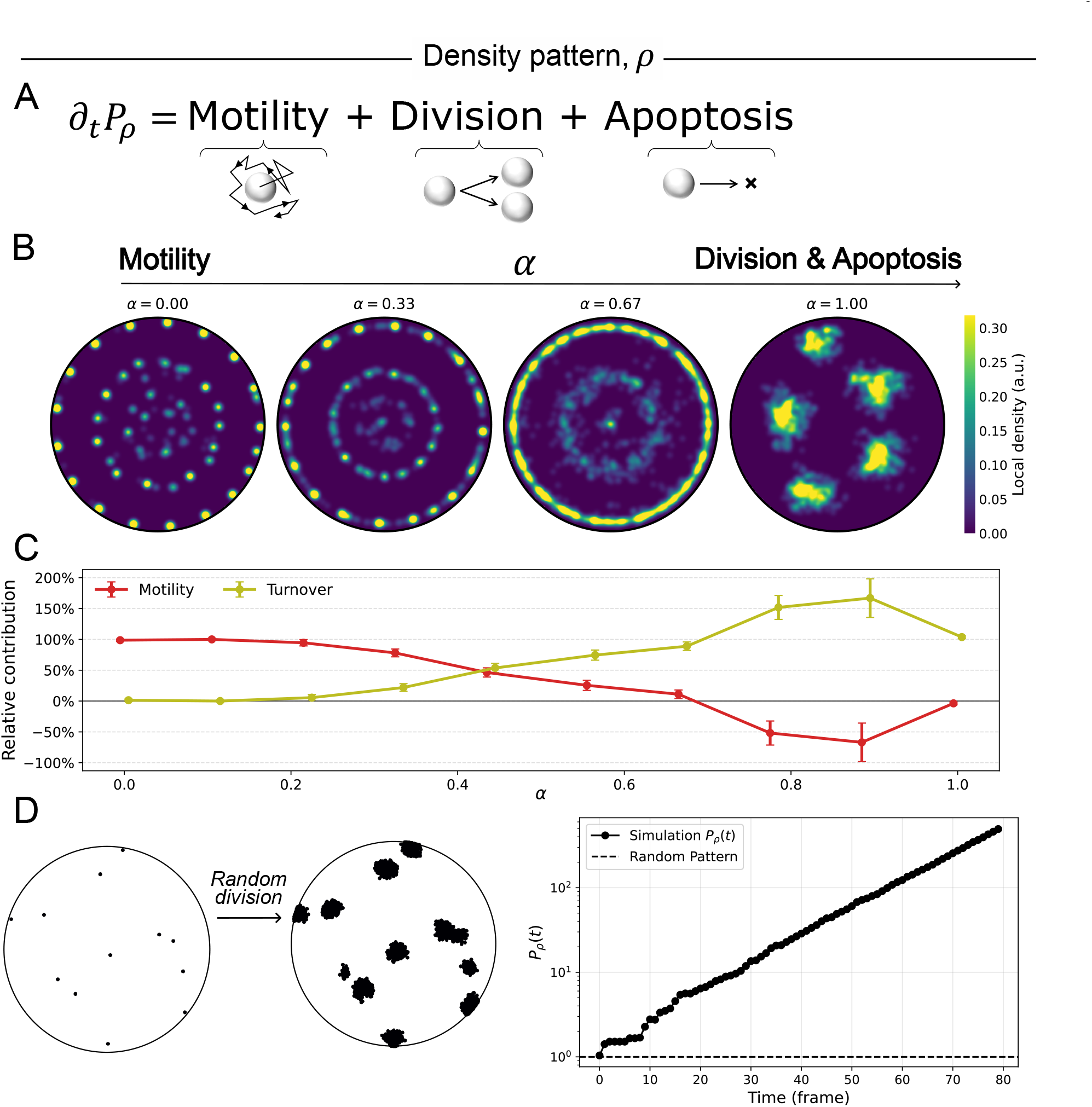
Validation of cellular contributions to cell number density pattern formation using agent-based models. (A) Exact decomposition of changes of the density pattern measure *P*_*ρ*_ into contributions by cell motility, cell division, and cell appearance/disappearance. (B) Snapshots of the agent-based model for different values of the mixing parameter *α*, tuning the relative contribution of cell motility and cell turnover, i.e. division and apoptosis, to pattern formation. (C) Relative contributions of motility and cell turnover to density pattern formation as a function of the mixing parameter *α*. Our decomposition correctly recovers the dominant pattern-forming process prescribed in the agent-based model. (D) Effect of cell divisions alone on density pattern formation. In the absence of motility and apoptosis, repeated cell divisions increase the density pattern measure *P*_*ρ*_.

**Figure 3.**
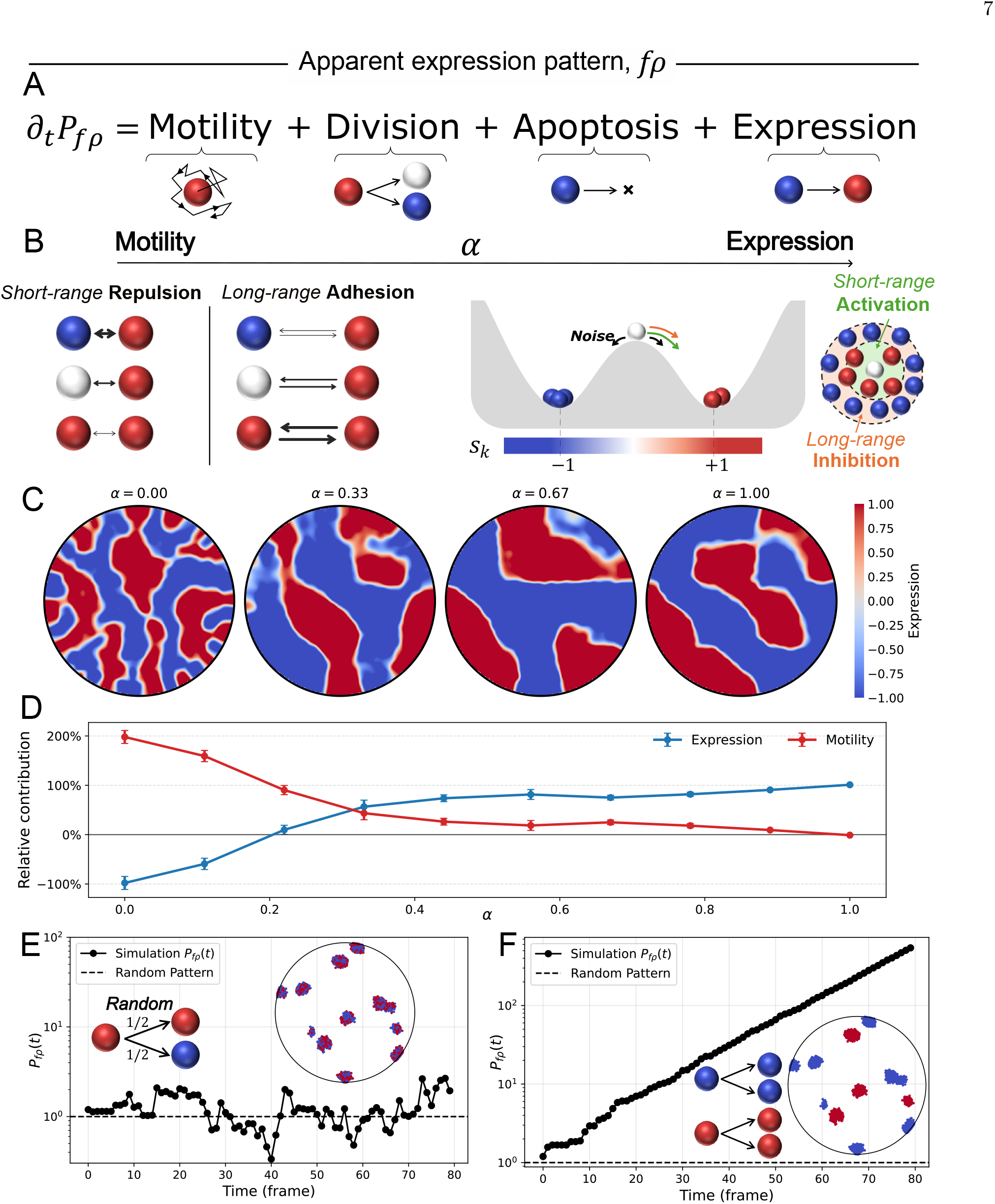
Validation of cellular contributions to apparent expression pattern formation using agent-based models. (A) Exact decomposition of changes in the apparent expression pattern measure *P*_*fρ*_ into contributions by cell motility, cell division, cell appearance/disappearance, and changes in gene expression. (B) Schematic illustration of the agent-based model combining cell motility and gene expression dynamics, with their relative contribution to pattern formation controlled by the tuning parameter *α*. (C) Snapshots of apparent expression patterns generated by the agent-based model for different values of the tuning parameter *α*. (D) Relative contributions of motility and gene expression dynamics to apparent expression pattern formation as a function of the tuning parameter *α*. The decomposition correctly identifies the dominant pattern-forming process prescribed in the agent-based model. (E) Effect of cell divisions, where each daughter cell acquires a random expression state. In this case, *P*_*fρ*_ remains close to the random-pattern baseline. (F) Effect of cell divisions with perfectly inherited expression states. This leads to a strong increase in the apparent expression pattern measure *P*_*fρ*_.

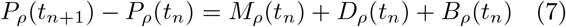

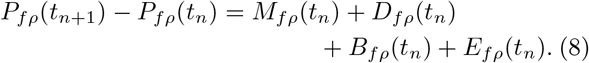

Here, *M*_*ρ/fρ*_, *D*_*ρ/fρ*_, *B*_*ρ/fρ*_, and *E*_*fρ*_ correspond to the respective contributions by cell motion, cell divisions, cell appearance & disappearance, and change of protein expression. The contribution *B*_*ρ/fρ*_ includes cell apoptosis events, but also cell appearance and disappearance across the boundary of the observation domain, as well as – in practice – gaps in the cell tracks. Our derivation also shows that there can be no other cellular contribution.

#### 2. Validation

We validate our framework using data simulated by agent-based models. Using such models, we explicitly prescribe the cellular processes driving pattern formation, and then test whether Eqs. (7) and (8) correctly identify them.

##### a. Cellular contributions to cell density pattern formation.

We test Eq. (7) decomposing the change of *P*_*ρ*_ into contributions by cell motility, cell division, and apoptosis (Figure 2A). We developed a minimal model that allows to tune which of these processes drives pattern formation through a parameter *α* (Figure 2B; see SI for full model details). For *α* = 0, pattern formation is exclusively driven by motility, while for *α* = 1, it is exclusively driven by cell turnover, i.e. cell division and apoptosis. A value of 0 *< α <* 1 indicates that pattern formation is driven by a combination of motility and turnover. Motility is modeled combining short-range repulsion, long-range adhesion, and Brownian motion, all restricted to a circular domain of radius *R*. For cell turnover we introduce asymmetric interaction ranges where division is locally activated via short-range interactions, whereas apoptosis is activated by long-range interactions. This asymmetry enables the emergence of spatial heterogeneity even in the absence of cell motility.

For all values of *α*, starting from a random uniform distribution of cells, we observed pattern formation both visually and through a temporal increase in *P*_*ρ*_ (Figure 2B). We then applied Eq. (7) to the simulated trajectories, and the obtained cellular contributions were fully consistent with what we prescribed in the model (Figure 2C): For *α* = 0, we find that the increase in *P*_*ρ*_ is entirely due to cell motility, *M*_*ρ*_. Conversely, for *α* = 1, the increase in *P*_*ρ*_ is almost entirely due to cell turnover, *D*_*ρ*_ and *B*_*ρ*_. For intermediate values of *α*, the method identifies mixed contributions from motility and turnover, where the turnover contribution generally increases with *α*. Note that in some of the mixed cases, contributions were smaller than 0% or larger than 100%. For instance at *α ≈* 0.9, pattern formation is driven by turnover, but cell motility creates noise that degrades the pattern. As a consequence, motility *negatively* contributes to *P*_*ρ*_, leading to a turnover contribution larger than 100%. Taken together, these results demonstrate that our framework faithfully recovers the cellular processes driving multi-cellular density pattern formation, without requiring any prior knowledge of the underlying cellular interaction rules.

Finally, we briefly test the effect of cell divisions alone. In the absence of any motility or apoptosis, cell division is expected to increase *P*_*ρ*_, since it increases the number of cells, and *P*_*ρ*_ is expected to scale linearly with *N*. This is indeed what we observe starting from a set of 12 initial cells (Figure 2D, compare also Figure 1D). Indeed, Eq. (7) correctly captures the cause of the increase in *P*_*ρ*_ fully through the division contribution *D*_*ρ*_.

##### b. Cellular contributions to apparent expression pattern formation.

We first test Eq. (8) for apparent expression pattern formation driven by the interplay between motility and gene expression. Similarly to the previous section, we use a parameter *α* to tune the model between purely motility-driven pattern formation (*α* = 0) and purely gene-expression-driven pattern formation (*α* = 1). Cell motility is implemented similar to the previous section, where here in addition the mechanical pair interactions are modulated by the expression levels of both cells (Figure 3B left). The gene expression dynamics for any cell *i* are modeled using a continuous variable *f*_*i*_, which is subject to two preferred states at values *−* 1 and +1, formally captured by a double-well landscape (Figure 3B right). We furthermore include noise to allow for stochastic switches between the two states. Finally, we include terms that tend to more strongly pull the value of *f*_*i*_ to one of the two preferred states, depending on the surrounding cells; these interactions introduce alignment with the expression state of closely neighboring cells, and misalignment with the expression state of cells that are further away (Figure 3B right). Full details are provided in the SI.

We find pattern formation for all probed values of *α*, both visually and through a temporal increase in *P*_*fρ*_ (Figure 3C). Applying our framework to simulated trajectories, we further observe that the inferred contributions accurately reflect the prescribed patterning mechanisms (Figure 3D): Motility-driven regimes are correctly identified when *α* is small, expression-driven regimes dominate when *α* approaches one, and mixed contributions are recovered at intermediate values. These results demonstrate that Eq. (8) can quantitatively disentangle the respective roles of motility and gene expression in pattern formation.

Finally, we test whether Eq. (8) correctly captures the effect of cell divisions on apparent expression pattern formation. To this end, we study two scenarios, both of which allow for two discrete expression states, *f*_*i*_ = *−*1 (blue) and *f*_*i*_ = +1 (red). First we study a model where after any cell division, each daughter acquires a random state *f*_*i*_ (Figure 3E). In this case, *P*_*fρ*_ does not increase but it remains close to one, indicating a random pattern. This is different when each daughter inherits the expression state from the mother (Figure 3F). In this case, even when starting from a random cell pattern, *P*_*fρ*_ increases exponentially, similar to Figure 2D. In both cases, Figure 3E,F, the division contribution *D*_*fρ*_ fully captures the observed changes in *P*_*fρ*_. Taken together, our results demonstrate that Eq. (8) reliably dissects the cellular contributions to apparent expression pattern formation.

## C. Application to experimental data

### 1. Presomitic mesoderm cell aggregates

We applied our framework to experimental data on quasi-2D aggregates of mouse presomitic mesoderm cells, which are constrained to a circular region [20, 48]. These systems display the emergence of patterns both in terms of cell number density (Figure 4A) and of the expression of the protein T/Brachyury (T/Bra; Figure 4D). An important question is how much different cellular processes contribute to these patterns. One possibility is that pattern formation is created mostly by cell motion, which could drive accumulation of cells in some clusters. Another possibility is the locally correlated change of T/Bra expression, driven for instance by signaling interactions, followed by a subsequent accumulation of cells in high-T/Bra-expression regions through, e.g., chemotactically driven cell motion, and/or biased division and apoptosis.

**Figure 4.**
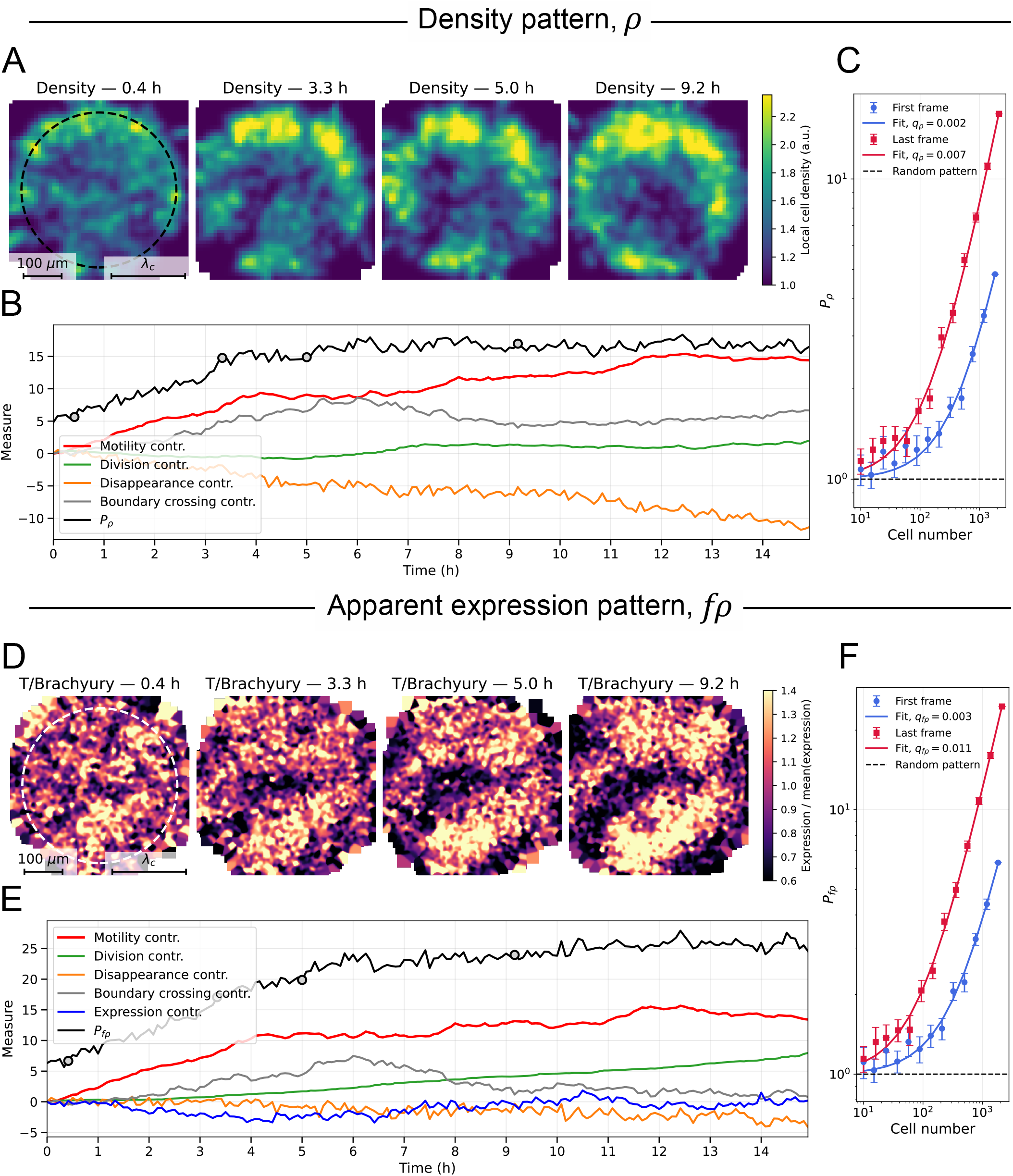
Pattern formation and cellular contributions in presomitic mesoderm cell aggregates. (A) Spatial distribution of cell density in the same aggregate at four time points (*t* = 0.4, 3.3, 5.0 and 9.2 h). Density maps are reconstructed from tracked cell positions and shown as smoothed spatial fields for visualization. (B) Temporal evolution of the density pattern measure *P*_*ρ*_(*t*) together with the decomposition of *P*_*ρ*_(*t*) *− P*_*ρ*_(0) into contributions by cell motility, cell division, boundary crossings, and cell disappearance. (C) Dependence of *P*_*ρ*_ on population size *N* obtained by subsampling, showing the temporal increase of the signal-to-noise ratio per cell. (D) Apparent T/Bra expression in the same aggregate at the same time points as in A, normalized by the spatial average. (E) Temporal evolution of the apparent expression pattern measure *P*_*fρ*_(*t*) together with the decomposition of *P*_*fρ*_(*t*) *− P*_*fρ*_(0) into contributions by cell motility, gene expression dynamics, cell division, boundary crossings, and cell disappearance. (F) Dependence of *P*_*fρ*_ on population size *N*, confirming the temporal increase of the signal-to-noise ratio per cell.

Recent work on the quasi-2D aggregates suggests that motion-driven cell aggregation precedes any patterning by changes in gene expression [20]. This would be partly consistent with observations in 3D mouse embryonic stem cell aggregates called gastruloids. These 3D aggregates usually give rise to a polarized T/Bra pattern, and recent work pointed to an important role of cell motion in driving pattern formation [19, 31]. However, other works on the 3D gastruloids highlight also an important role for cell differentiation in patterning [49, 50].

We applied our geometric formalism to quantify the cellular contributions to patterning in the quasi-2D aggregates. Experimentally, the cells are confined to a circular region, and for our analysis we use a circular domain. However, the confinement is not perfect, and many cells still cross the boundary of our circular analysis domain. We thus further divided the dis-/appearance contribution *B*_*ρ*_ into two contributions: one by crossing of the circular boundary of the analysis domain and one by cell disappearance (see SI section 5).

We first analyzed pattern emergence at the level of cell density, and then extended the analysis to spatial patterns of T/Bra expression. We find that the cell density pattern measure *P*_*ρ*_(*t*) increases during the experiment and subsequently reaches a plateau (Figure 4A, black curve in B). To identify the processes responsible for this increase, we decomposed the temporal variation of *P*_*ρ*_(*t*) into cellular contributions. We find that density pattern formation is driven by cell motility, while the contribution by divisions is small (Figure 4B). This is also consistent with the observation that not only *P*_*ρ*_(*t*) increased, but also the signal-to-noise ratio per cell, *q*_*ρ*_, (Figure 4C), indicating that the increase in *P*_*ρ*_(*t*) is not merely driven by an increase in cell number (as opposed to Figure 2D). Boundary crossings and cell disappearance provide small contributions, which almost cancel each other out. Taken together, these results indicate that density patterns in these aggregates primarily emerge from cell motion rather than cell turnover.

We next analyzed the apparent T/Bra expression pattern using the measure *P*_*fρ*_(*t*). We find an increase over time with a plateau starting at ~ 6*h* (Figure 4D, black curve in E, F). Decomposition into cellular contributions shows that pattern formation is again predominantly driven by cell motility (red curve in Figure 4E). In contrast, contributions from cellular T/Bra expression changes, divisions, boundary crossings, and disappearances play only a minor role (Figure 4E). That the mere change of cell number through divisions does not play a major role is also confirmed by the increase of the signal-to-noise ratio per cell (Figure 4F). This indicates that the apparent T/Bra expression pattern largely emerges from cellular motion rather than coordinated T/Bra expression.

Taken together, we found that almost exclusively cell motility is responsible for the emergence of both density and apparent T/Bra expression patterns. Similar results are obtained for a second data set (Supplementary Fig. S4). While these findings are consistent with earlier results [20], we stress that (i) our method requires only the cell tracks, and no additional perturbation experiment, and (ii) our purely geometric approach is completely independent of any mechanistic assumptions.

### 2. Breast cancer cell populations

As a second example, we applied our framework to published experimental data on 2D cultures of breast cancer cells [24]. These cultures contain different cell types, including a population enriched in cancer stem cells (CSCs) and differentiated cells, where the CSC-enriched population is marked by the expression of the ALDH1A1 protein [24, 51]. Expression patterns formed in these experiments are akin to stem cell niches, and understanding their emergence is important to tackle, for instance, immune escape and cancer treatment resistance. Here, we analyze the emergence of both cell number density patterns and apparent ALDH1A1 expression patterns.

In this example, the cells are experimentally not constrained within a circular region. To still apply our method, we select as region of interest the largest circle that can be inscribed into the field of view. Again, the effect of the cells that enter and leave this circle are explicitly accounted through the boundary crossing contribution (see SI section 5).

We first analyzed the formation of the density pattern. We observe the progressive emergence of large-scale density patterns (Figure 5A), where the measure *P*_*ρ*_(*t*) increases over time (Figure 5B, black curve). Decomposition into cellular contributions shows that pattern formation is primarily driven by cell division with a small contribution by cell dis-/appearance (Figure 5B). In contrast, the contributions by cell motility and boundary crossings are small. Cell division and dis-/appearance do not act merely through net increase in cell number. Instead, the signal-to-noise ratio per cell does also increase from *q*_*ρ*_ = 0.005 to *q*_*ρ*_ = 0.009 (Figure 5C), suggesting that these processes are biased to sharpen the pattern. Indeed, explicitly testing this hypothesis, we find that cells were more likely to divide in denser regions, at least at late times (Supplementary Fig. S5).

**Figure 5.**
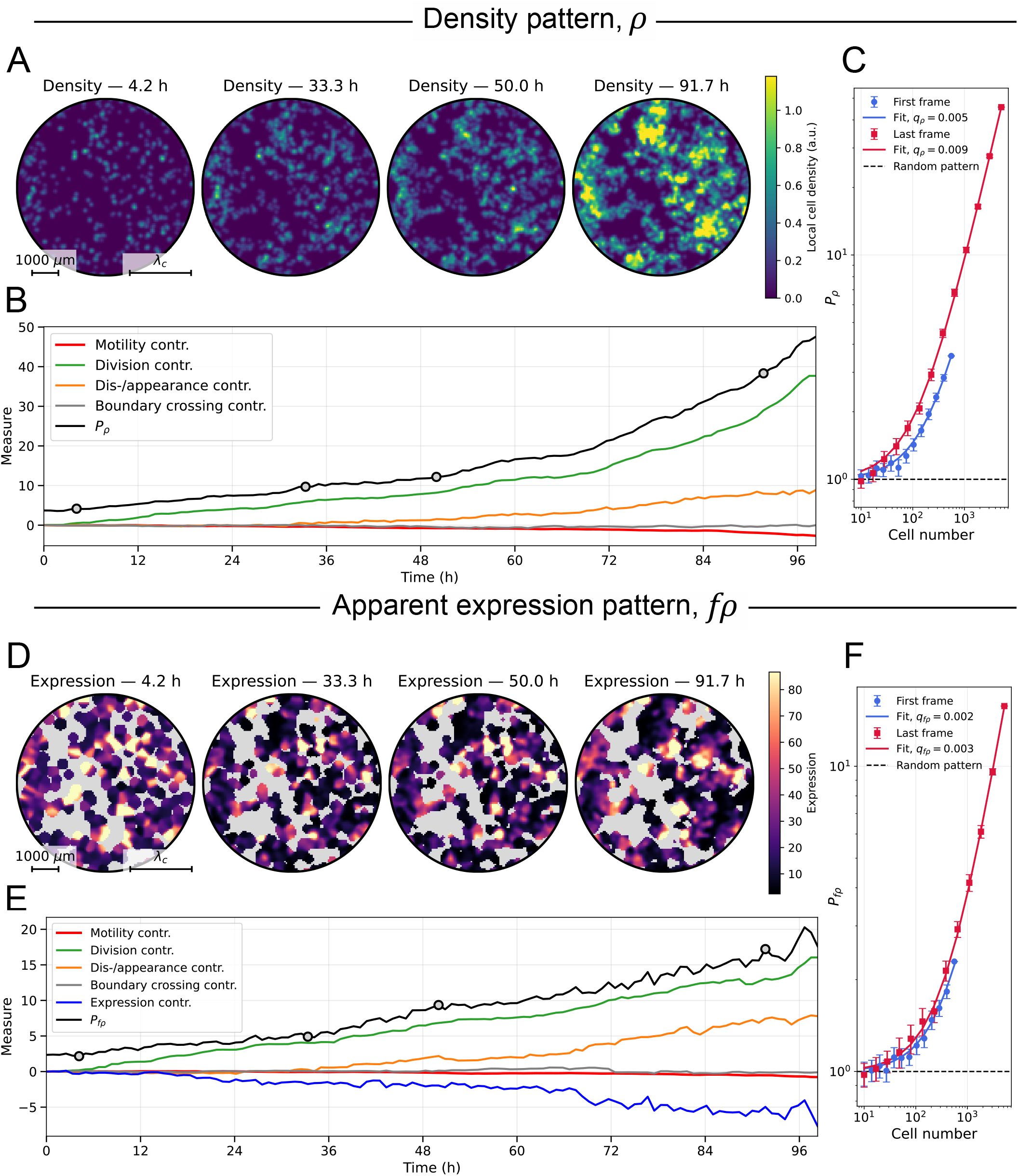
Pattern formation and cellular contributions in breast cancer cell populations. (A) Spatial distribution of cell density at four time points (*t* = 4.2, 33.3, 50.0 and 91.7 h). Density maps are reconstructed from tracked cell positions and shown as smoothed spatial fields for visualization. (B) Temporal evolution of the density pattern measure *P*_*ρ*_(*t*) together with the decomposition of *P*_*ρ*_(*t*) *− P*_*ρ*_(0) into contributions by cell motility, cell division, boundary crossings, and cell dis-/appearance. (C) Dependence of *P*_*ρ*_ on population size *N* obtained by subsampling, showing the temporal increase of the signal-to-noise ratio per cell. (D) Apparent ALDH1A1 expression pattern at the same time points as in A, shown after normalization by the spatial average. Areas devoid of cells are shown in gray. (E) Temporal evolution of the apparent expression pattern measure *P*_*fρ*_(*t*) together with the decomposition of *P*_*fρ*_(*t*) *− P*_*fρ*_(0) into contributions by cell motility, gene expression dynamics, cell division, boundary crossings, and cell dis-/appearance. (F) Dependence of *P*_*fρ*_ on population size *N*, showing that the signal-to-noise ratio per cell remains approximately constant during the observation window.

We next examined the dynamics of the apparent ALDH1A1 expression pattern using *P*_*fρ*_(*t*) (Figure 5D). The measure *P*_*fρ*_(*t*) increases over time (Figure 5E, black curve), in a way mostly driven by cell divisions (Figure 5E, green curve). However, the signal-to-noise ratio per cell increases only little, from *q*_*fρ*_ *≈* 0.0022 to *q*_*fρ*_ *≈* 0.0028 (Figure 5F). This indicates that the strong temporal increase of *P*_*fρ*_(*t*) is mostly driven by an increase in cell number. In contrast, the cell motility and boundary crossing contributions are essentially zero. We further find that changes in ALDH1A1 expression contribute negatively to pattern formation, an effect that is over-compensated by both cell divisions and cell dis-/appearance (Figure 5E). One possible reason for this is that while inheritance of transcriptional cell states across several divisions has been observed before, that inheritance is imperfect and gets lost over time [24] (compare to perfect cell state inheritance in Figure 3F).

Taken together, we find that pattern formation is dominated by cell divisions, which is further confirmed by the analysis of a second data set (Supplementary Fig. S6). Divisions create cell number density patterns, where an increase in the signal-to-noise ratio indicates that cell divisions are biased towards denser regions. The apparent “stem cell niche” patterns evolve through an interaction between cell divisions and expression. Earlier work suggested that both cell division and expression changes contributed positively to ALDH1A1 pattern formation, while cell motility contributed negatively [24]. However, by directly connecting cellular processes to their effect on large-scale patterns, our method revealed instead that the net effect of expression changes acts to *deteriorate* the ALDH1A1 pattern, e.g. due to the loss of inherited cell state over time. Meanwhile, cell motility has essentially no effect on pattern formation in this system, neither positive nor negative.

## III. DISCUSSION

We proposed a method to quantify multi-cellular pattern formation, and the cellular processes driving it. Specifically, we introduced measures for cell number density patterns, *P*_*ρ*_ and apparent expression patterns, *P*_*fρ*_, and we decomposed them into contributions by cell expression changes, cell motility, cell divisions, and cell apoptosis. To automatize computation, we developed the publicly available python tool CREDIT (Cellular Roles in Emergent Development Inferred from Tracks).

We used a very generic definition for a pattern: any fluctuation that occurs above some cutoff wavelength *λ*_*c*_. This notably also includes non-periodic patterns. Our approach is purely geometric and does not require any assumptions about the mechanism driving pattern formation, it only requires the information about cell tracks and expression levels. In other words, our method quantifies *what* the cells are doing that drives pattern formation, without asking *why* they are doing it. Our work provides the foundation for subsequent hypothesis-driven work, using for instance mechanistic models.

Using simulated data, we validated important features of our method, which are: (i) The pattern measures are normalized such that their value directly provides the significance level for whether there is a pattern (Figure 1C inset). Note that our normalization automatically compensates for effects such as photobleaching. (ii) In the presence of a – potentially noisy – pattern, our pattern measures scale with the number of cells *N* (Figure 1D,F). Correspondingly, when changing the cell number for the same pattern and noise level, the signal-to-noise ratios per cell, *q*_*ρ*_ and *q*_*fρ*_, remain the same. (iii) The pattern measures depend on a cutoff length scale *λ*_*c*_ that separates what shall be interpreted as small-scale fluctuations from large-scale pattern (Figure 1E). (iv) We also validated the decomposition into cellular contributions for different scenarios (Figure 2, Figure 3).

We further demonstrated our approach on experimental data on both quasi-2D aggregates of mouse embryonic cells [20] and on 2D colonies of cancer cells [24]. For the quasi-2D aggregates of mouse embryonic cells, our method shows that the formation of both cell density and apparent expression patterns are mostly driven by cell motion. This highlights the central role of mechanical interactions driving coordinated cell movements in this system [20]. For the 2D colonies of cancer cells, our method shows that density and apparent expression pattern formation is driven mostly by cell divisions. Based on cell lineage analysis, previous work additionally suggested a positive contribution of gene expression to pattern formation [24]. However, linking single cell trajectories and lineage to the global pattern, our method revealed that, on the contrary, the net contribution by gene expression level changes is negative. This was revealed thanks to the fact that our geometric approach directly and quantitatively links cell-scale processes to the large-scale pattern formation.

Our method can be applied to both experimental and simulated data. For our validations in Figures 2 and 3, we used a computational model where the processes driving pattern formation are clearly defined. However, our method will also be important in theoretically studying the behavior of agent-based models where the cellular processes driving pattern formation are less clear [52–55].

A number of variations of our approach are possible. First, here, we focused on a circular region of interest, and correspondingly used a decomposition based on Bessel-Fourier modes. However, in particular for simulations of agent-based models with periodic boundary conditions, we recommend instead usual Fourier modes. More generally, for arbitrarily shaped domains, one could instead use, e.g., the eigen functions of the Laplace operator on that domain. Second, here, we demonstrated the application to 2D systems, but our approach can be directly generalized to 3D systems. Third, in some situations, it can make sense to additionally modify the cellular expression levels *f*_*i*_. For instance, if the average of the *f*_*i*_ is much larger than their standard deviation, the apparent expression pattern is almost the same as the density pattern. In such cases, it could be useful to instead use the expression levels corrected by their average *f*_*i*_ *−⟨f⟩* (see SI section 5). Fourth, in the future, it could be interesting to develop a pattern measure for the expression pattern alone, *f*, instead of the apparent expression pattern, *fρ* (compare Figure 1A). Fifth, our method can in principle also generalized to patterns consisting of multiple gene expression markers, which we will consider in future work. Finally, it would be interesting to develop a decomposition of existing pattern measures, e.g. correlation functions [42, 43], into cellular contributions and compare to the method presented here.

By providing a quantitative, model-free decomposition of pattern formation, our framework bridges the gap between single-cell dynamics and collective cellular organization. As imaging and tracking technologies continue to advance, in particular in 3D systems, such datadriven approaches are likely to play an increasingly important role in uncovering the dynamical principles of self-organization across biological systems.

## ACKNOWLEDGMENTS

We thank Jan Rombouts, Anna Erzberger, François Anquez, Chann Lagadec, and Benjamin Pfeuty for useful discussions. This project received funding from the European Research Council (ERC) under the European Union’s Horizon Europe research and innovation program – ERC Synergy Grant 101072123. A CC-BY public copyright license has been applied by the authors to the present document and will be applied to all subsequent versions up to the Author Accepted Manuscript arising from this submission, in accordance with the grant’s open access conditions.

