## Supplemental Information for "Quantification of cellular contributions to multi-cellular pattern formation^*^"

### Supplementary Methods : Quantification of cellular contributions to multi-cellular pattern formation

October 6, 2026

**10 Testing for density-dependent cell division in breast cancer cell populations** **29**

**11 Code availability** **30**

### 1 Fourier–Bessel decomposition: mathematical definition

We consider a set of tracked cells characterized at each time point by their positions $\mathbf{x}_k = (r_k, \theta_k)$ in polar coordinates and by an associated scalar quantity $f_k$. All spectral analyses are performed on a circular domain of fixed radius $R$.

#### 1.1 Fourier–Bessel basis

To decompose scalar fields defined on a disk, we use the Fourier–Bessel eigenfunctions of the Laplacian subject to homogeneous Neumann boundary conditions. The complete Neumann eigenbasis contains a constant mode, $\phi_{0,0}(r,\theta) = 1$, which is associated with the zero eigenvalue of the Laplacian.

The nonconstant modes are indexed by $z = (m, n)$, with $m \in \mathbb{Z}$ and $n \geq 1$, and are defined as

$$\phi_{m,n}(r,\theta) = J_{|m|}\left(\beta_{|m|,n}\frac{r}{R}\right) e^{im\theta}.$$

Here, $J_m$ denotes the Bessel function of the first kind of order $m$, and $\beta_{m,n}$ is the $n$-th strictly positive zero of $J'_m$, i.e.

$$J'_m(\beta_{m,n}) = 0, \qquad \beta_{m,n} > 0.$$

These modes satisfy the Laplacian eigenvalue problem $-\Delta\phi_{m,n} = k^2_{m,n}\phi_{m,n}$, where $k_{m,n} = \beta_{|m|,n}/R$ is the associated wavenumber. We also define the corresponding characteristic length scale as

$$\lambda_{m,n} = \frac{2\pi}{k_{m,n}} = \frac{2\pi R}{\beta_{|m|,n}}, \qquad (m,n) \in \mathcal{Z}^*.$$

For the constant mode, $k_{0,0} = 0$, corresponding formally to an infinite characteristic length scale.

For later use, we denote the complete index set by $\mathcal{Z} = \{(0,0)\} \cup \{(m,n) : m \in \mathbb{Z},\ n \geq 1\}$, and the set of nonconstant modes by $\mathcal{Z}^* = \mathcal{Z} \backslash \{(0,0)\}$. Although the constant mode belongs to the complete Neumann eigenbasis, it is excluded from the spectral pattern measures introduced below because it represents only the spatially homogeneous component of the field.

#### 1.2 Normalization of the modes

The squared norm of each Fourier–Bessel mode is defined as

$$K_{m,n} = \int_0^{2\pi}\int_0^R |\phi_{m,n}(r,\theta)|^2 \, r \, dr \, d\theta.$$

For the constant mode, the normalization is

$$K_{0,0} = \pi R^2.$$

For the nonconstant radial modes, the normalization is

$$K_{0,n} = \pi R^2 J_0^2(\beta_{0,n}), \qquad n \geq 1.$$

For the nonconstant angular modes, the normalization is

$$K_{m,n} = \pi R^2 \left(1 - \frac{|m|^2}{\beta^2_{|m|,n}}\right) J^2_{|m|}(\beta_{|m|,n}), \qquad |m| \geq 1, \quad n \geq 1.$$

##### 1.3 Definition of the spectral coefficients

Given a scalar field $f(r,\theta)$ defined on the disk, the corresponding Fourier–Bessel coefficients are defined by projection onto the normalized modes associated with the orthogonal functions $\phi_z$. For the constant mode, one has

$$d_{0,0} = \frac{1}{\sqrt{K_{0,0}}} \int_0^{2\pi} \int_0^R f(r,\theta)\, r\, dr\, d\theta.$$

For each nonconstant mode $z = (m,n) \in \mathcal{Z}^*$, the coefficient is

$$d_{m,n} = \frac{1}{\sqrt{K_{m,n}}} \int_0^{2\pi} \int_0^R f(r,\theta) J_{|m|}\Big(\beta_{|m|,n} \frac{r}{R}\Big)\, e^{-im\theta}\, r\, dr\, d\theta.$$

For real-valued fields, the coefficients satisfy

$$d_{-m,n} = \overline{d_{m,n}}.$$

The field can then be reconstructed as

$$f(r,\theta) = \frac{d_{0,0}}{\sqrt{K_{0,0}}} + \sum_{n=1}^{\infty} \frac{d_{0,n}}{\sqrt{K_{0,n}}} J_0\Big(\beta_{0,n} \frac{r}{R}\Big) + 2 \sum_{m=1}^{\infty} \sum_{n=1}^{\infty} \Re\left[ \frac{d_{m,n}}{\sqrt{K_{m,n}}} J_m\Big(\beta_{m,n} \frac{r}{R}\Big)\, e^{im\theta} \right].$$

Since $K_{0,0} = \pi R^2$, the constant contribution is

$$\frac{d_{0,0}}{\sqrt{K_{0,0}}} = \frac{1}{\pi R^2} \int_0^{2\pi} \int_0^R f(r,\theta)\, r\, dr\, d\theta,$$

which corresponds to the spatial average of the field over the disk.

##### 1.4 Particle-based approximation

In practice, the scalar field is represented as a distribution-valued field composed of a discrete set of weighted point-like elements corresponding to tracked cells. Denoting by $\Omega(t)$ the set of cells present at time $t$, we write

$$f(\mathbf{x}, t) = \sum_{k \in \Omega(t)} f_k(t)\, \delta(\mathbf{x} - \mathbf{x}_k(t)).$$

At discrete time points $t_i$, the corresponding nonconstant Fourier–Bessel coefficients are approximated by the particle-based expression

$$d_z(t_i) = \frac{1}{\sqrt{K_z}} \sum_{k \in \Omega(t_i)} f_k(t_i)\, J_{|m|}\left(\beta_{|m|,n} \frac{r_k(t_i)}{R}\right) e^{-im\theta_k(t_i)}, \qquad z \in \mathcal{Z}^*.$$

This Dirac-based approximation preserves the spectral structure of the continuous decomposition while allowing direct application to single-cell tracking data. All pattern measures and temporal decompositions used in this work are constructed from these coefficients.

### 2 Null distribution of the spectral pattern measure

#### 2.1 Statistical baseline and assumptions

To assess whether an observed value of the spectral pattern measure reflects genuine spatial structure or can arise from spatially uncorrelated variability, we introduce a statistical baseline describing a spatially unstructured system. Even in the absence of spatial patterning, finite sampling effects generically produce nonzero contributions to low-wavenumber modes.

We consider a simple reference model in which cell positions are randomly and uniformly distributed over a disk of radius $R$, and the cell-associated scalar quantity does not depend on position. Denoting by $\mathbf{x}_k = (r_k, \theta_k)$ the position of cell $k$ and by $f_k$ the associated scalar value, the baseline assumptions are

$$\mathbf{x}_k \sim \mathrm{Unif}(\mathcal{D}(R)), \qquad f_k \text{ i.i.d. with mean } \mu_f \text{ and variance } \sigma_f^2, \quad \text{independent of } \mathbf{x}_k.$$

For experimental data, the values of $\mu_f$ and $\sigma_f^2$ are computed from the final time point of each experiment, which is the stage at which spatial patterning is evaluated. This construction preserves the overall distribution of the measured signal while removing any spatial relationship between signal values and cell positions.

Under these assumptions, any apparent large-scale pattern arises solely from random spatial sampling and finite population size, rather than from structured spatial coupling between positions and scalar values. Deviations from this baseline therefore provide a quantitative criterion to assess the presence of statistically significant spatial patterning.

#### 2.2 Asymptotic distribution of the pattern measure

Under the statistical baseline introduced above, the pattern measure can be analytically approximated in the large-sample limit. In the following, we derive the null distribution for the apparent expression pattern measure $P_{f\rho}$, where each cell carries a scalar value $f_k$ independent of its position. The density pattern measure $P_\rho$ follows as the special case $f_k = 1$ for all cells.

**Asymptotic distribution of spectral coefficients.** For uniformly distributed cell positions and scalar values $f_k$ independent of position, each nonconstant Fourier–Bessel coefficient

$$d_z = \frac{1}{\sqrt{K_z}} \sum_{k=1}^{N} f_k J_{|m|}\left(\tfrac{\beta_{|m|,n} r_k}{R}\right) \mathrm{e}^{-im\theta_k}, \qquad z = (m,n) \in \mathcal{Z}^*.$$

is a sum of independent random variables with zero mean. By the central limit theorem, $d_z$ converges asymptotically to a Gaussian random variable. Modes with $m = 0$ are real-valued and normally distributed, whereas modes with $m \neq 0$ are proper circular complex Gaussians.

The variance of each mode can be computed explicitly and reads

$$\sigma_z^2 = \frac{N}{\pi R^2} (\mu_f^2 + \sigma_f^2) = \bar{\rho}\, \mathbb{E}[f^2] := \sigma^2, \qquad z \in \mathcal{Z}^*,$$

where $\bar{\rho} = N/(\pi R^2)$ denotes the mean cell density in the observation domain and $\mathbb{E}[f^2] = \mu_f^2 + \sigma_f^2$ denotes the second moment of the scalar quantity.

**Modal energies.** As a consequence, the modal energies $|d_z|^2$ follow asymptotically Gamma distributions. More precisely, for $m \neq 0$, $|d_z|^2/\sigma^2$ is exponentially distributed, while for $m = 0$ it follows a $\chi^2$ distribution with one degree of freedom.

**Spectral energy sums.** Let

$$S_{\mathrm{num}} = \sum_{\substack{n\geq 1\\ \lambda_{0,n}>\lambda_c}} |d_{0,n}|^2 + 2 \sum_{\substack{m\geq 1,\ n\geq 1\\ \lambda_{m,n}>\lambda_c}} |d_{m,n}|^2.$$

The first sum contains the retained nonconstant radial modes, whereas the second sum contains one representative of each conjugate pair $(m, -m)$. The factor of 2 accounts for the two real degrees of freedom associated with each pair.

Using the results above, each term in the first sum follows a Gamma distribution with shape parameter $1/2$ and scale parameter $2\sigma^2$, whereas each term in the second sum follows a Gamma distribution with shape parameter 1 and the same scale parameter $2\sigma^2$.

Since sums of independent Gamma variables with a common scale parameter remain Gamma-distributed, it follows that

$$S_{\mathrm{num}} \sim \Gamma\left(N_0/2 + N_+,\, 2\sigma^2\right),$$

with $N_0$ the number of retained nonconstant modes with $m = 0$ and $N_+$ the number of retained modes with $m > 0$.

Equivalently, introducing the total number of spectral degrees of freedom, $M_c = N_0 + 2N_+$, one obtains

$$S_{\mathrm{num}} \sim \Gamma\left(\frac{M_c}{2},\, 2\sigma^2\right).$$

**Distribution of the measure.** For clarity, we distinguish between a theoretical version of the measure, defined using the true second-order moment of the scalar quantity, and its empirical estimator used in practice.

The theoretical measure is defined as

$$P^{\star}_{f\rho} = \frac{S_{\mathrm{num}}}{M_c\,\bar{\rho}\,\mathbb{E}[f^2]}.$$

Under the statistical baseline, the measure $P^{\star}_{f\rho}$ is a Gamma random variable, given by

$$P^{\star}_{f\rho} \sim \Gamma\left(\frac{M_c}{2},\, \frac{2}{M_c}\right),$$

with mean $\mathbb{E}[P^{\star}_{f\rho}] = 1$ and variance $\mathrm{Var}(P^{\star}_{f\rho}) = 2/M_c$ .

In practice, the second-order moment $\mathbb{E}[f^2]$ is unknown and is replaced by its empirical estimator

$$\overline{f^2} = \frac{1}{N}\sum_{k=1}^{N} f_k^2,$$

leading to the empirical pattern measure

$$\widehat{P_{f\rho}} = \frac{S_{\mathrm{num}}}{M_c\,\bar{\rho}\,\overline{f^2}}.$$

The use of $\overline{f^2}$ introduces additional fluctuations in the denominator, so that $\widehat{P_{f\rho}}$ is no longer exactly Gamma-distributed. However, in the large-sample limit, $\overline{f^2}$ converges to $\mathbb{E}[f^2]$. As a result, $\widehat{P_{f\rho}}$ remains well approximated by the same rescaled Gamma distribution.

**Monte Carlo validation of the null distribution.** The analytical prediction was tested by Monte Carlo simulations for both measures. The corresponding validation for the density pattern measure $P_\rho$ is shown in the main text, while the validation for the apparent expression pattern measure $P_{f\rho}$ is shown in Fig. S1. In both cases, the simulated null distributions are centered around $P = 1$ and agree with the predicted rescaled Gamma distribution.

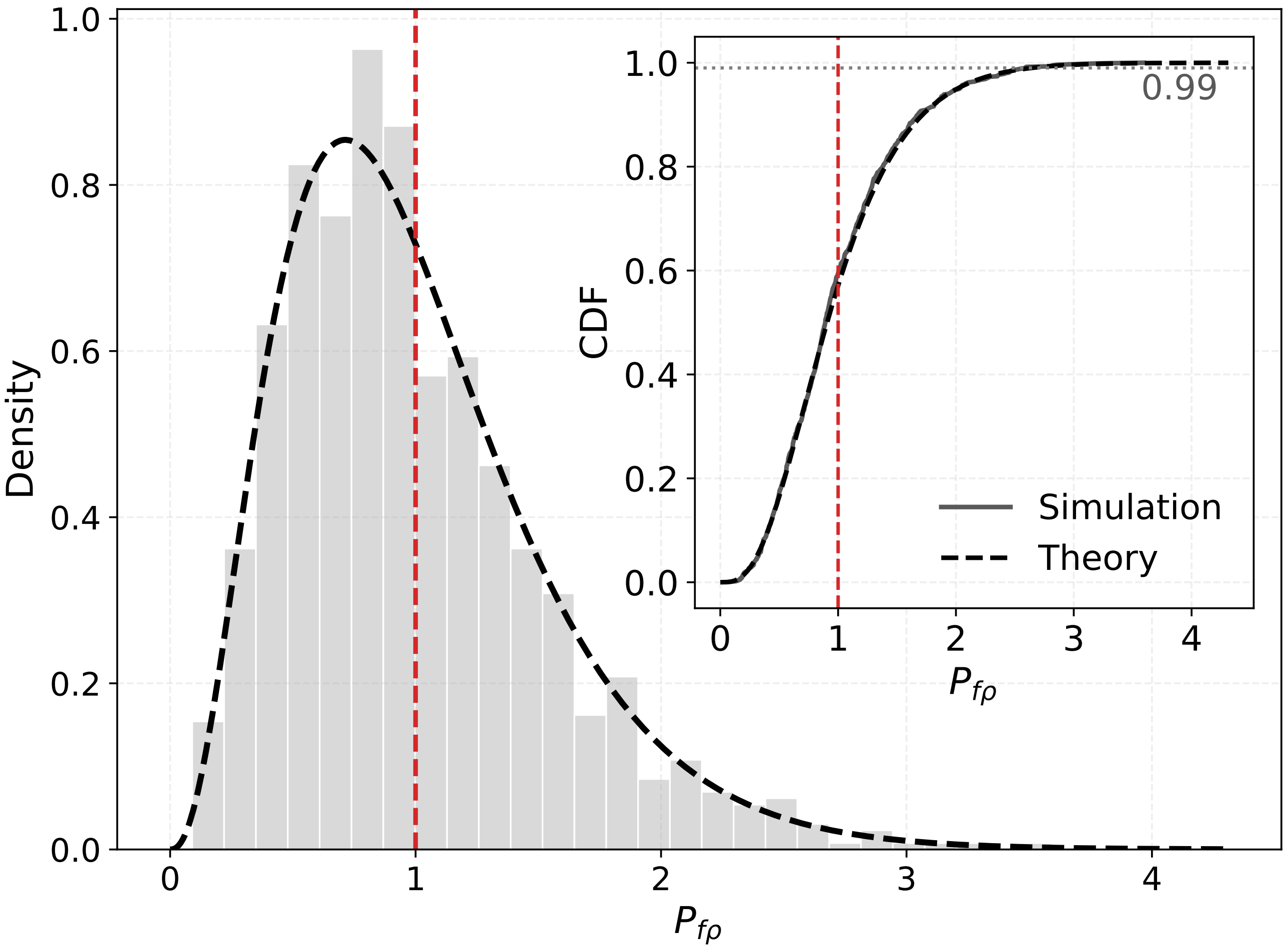

Figure S1: Null distribution of the apparent expression pattern measure $P_{f\rho}$. Cell positions were sampled uniformly in a disk, and expression values were assigned independently of cell position. Gray bars show the distribution obtained from Monte Carlo simulations, while the black dashed curve shows the analytical prediction. The red dashed line indicates the null expectation $P_{f\rho} = 1$.

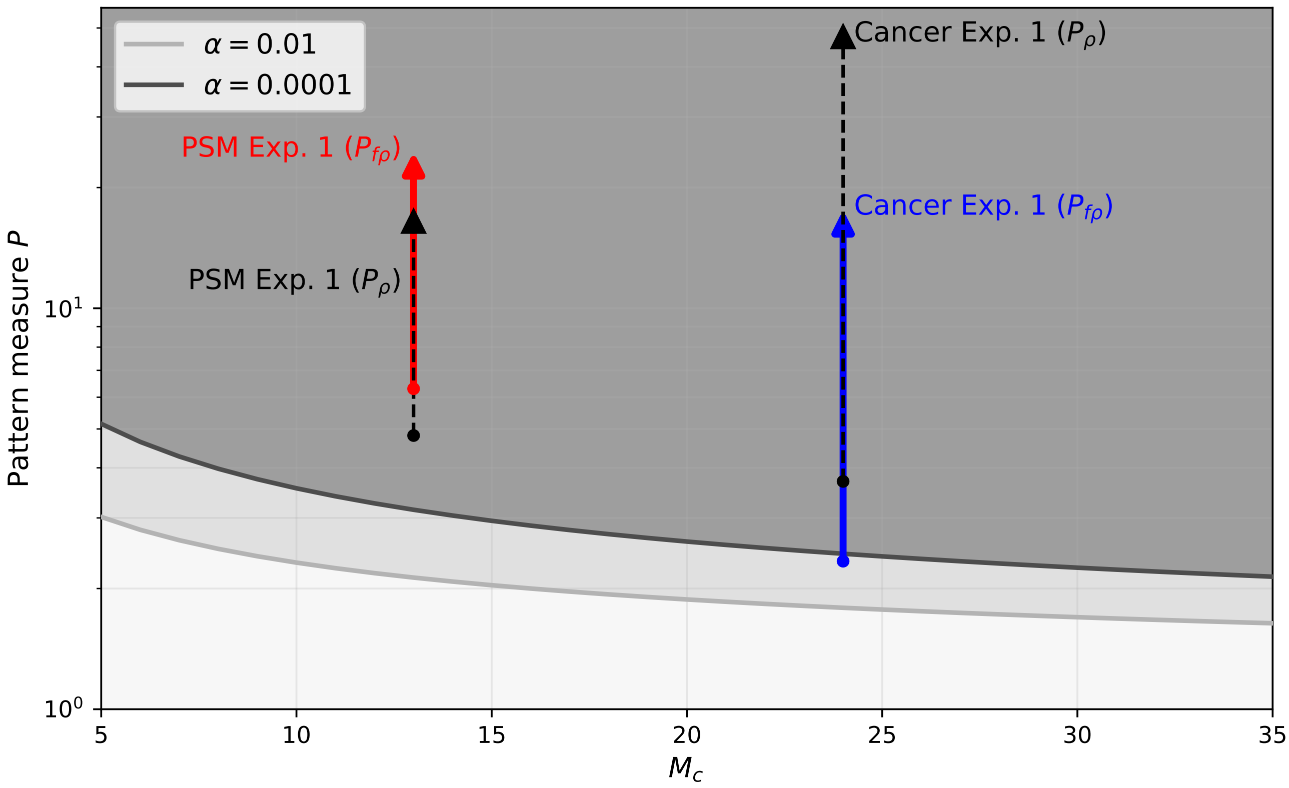

Figure S2: Statistical assessment of experimental pattern measures. Gray curves show significance thresholds for two values of $\alpha$, as a function of the number of nonconstant Fourier–Bessel modes $M_c$. Arrows indicate the change in $P$ from the initial to the final time point for $P_\rho$ and $P_{f\rho}$ in the presomitic mesoderm aggregate and breast cancer datasets. Values above a threshold correspond to statistically significant patterns at the corresponding significance level. Here, $M_c = 7$ modes were taken into account to compute $P_{f\rho}$.

#### 2.3 Practical use for experimental data

For each experimental condition, the relevant pattern measure, either $P_\rho$ or $P_{f\rho}$, can be compared to the null distribution derived above. Given an observed value $P_{\text{obs}}$, we define the p-value as the upper-tail probability

$$p = \mathbb{P}_0(P \geq P_{\text{obs}}),$$

where $\mathbb{P}_0$ denotes the probability under the spatially unstructured null model.

Equivalently, for a chosen significance level $\alpha$, one can define a critical value $P_\alpha$ such that

$$\mathbb{P}_0(P \geq P_\alpha) = \alpha.$$

Observed values satisfying $P_{\text{obs}} \geq P_\alpha$ are therefore considered statistically significant.

In the experimental applications, we compare both the initial and final values of the pattern measure to this null distribution. This allows us to distinguish cases where a pattern is already present at the beginning of the movie from cases where the pattern measure crosses the significance threshold during the experiment.

In Fig. S2, we represent this statistical test by plotting the critical values $P_\alpha$ as a function of the number of retained Fourier–Bessel modes $M_c$. The arrows indicate the change in $P$ between the initial and final time points for each experimental case.

### 3 Subsampling method

#### 3.1 Scaling of pattern measures with cell number

The dependence of the pattern measures on the number of cells provides a way to distinguish random finite-sampling fluctuations from genuine spatial patterns. We summarize here the scaling argument underlying Eqs. (5) and (6) of the main text.

Consider a nonconstant spectral coefficient, either $c_z$ for the density pattern or $d_z$ for the apparent expression pattern, with $z \in \mathcal{Z}^*$. In both cases, the coefficient can be viewed as a sum of contributions from individual cells. In the absence of a spatial pattern, these contributions add incoherently. The mean contribution to a given large-scale nonconstant mode is then zero, while the variance of the sum increases linearly with the number of sampled cells. Therefore,

$$\mathbb{E}\left[|c_z|^2\right] \sim N, \qquad \mathbb{E}\left[|d_z|^2\right] \sim N.$$

Since the pattern measures are normalized by the mean cell density, which scales as $\bar{\rho} \sim N$, random configurations yield values of order one,

$$P_\rho(N) \sim 1, \qquad P_{f\rho}(N) \sim 1.$$

In contrast, when a spatial pattern is present, the contributions to the modes associated with that pattern have a nonzero average. For these modes, the spectral coefficient contains a mean component that scales as $N$. Consequently, the corresponding modal energy contains both a signal contribution scaling as $N^2$ and a finite-sampling contribution scaling as $N$,

$$\mathbb{E}\left[|c_z|^2\right] \sim N^2 + N, \qquad \mathbb{E}\left[|d_z|^2\right] \sim N^2 + N.$$

After normalization by $\bar{\rho} \sim N$, this gives a linear dependence of the pattern measure on cell number,

$$P_\rho(N) = 1 + q_\rho N, \qquad P_{f\rho}(N) = 1 + q_{f\rho} N.$$

The prefactors $q_\rho$ and $q_{f\rho}$ quantify the signal-to-noise ratio per cell for the density and apparent expression patterns, respectively.

#### 3.2 Random subsampling procedure

To estimate $q_\rho$ and $q_{f\rho}$ from simulated or experimental data, we used a random subsampling procedure. Starting from a configuration containing $N_{\text{tot}}$ cells, we selected random subsets of size $N$, with $N < N_{\text{tot}}$. For each subset, we recomputed the corresponding pattern measure, $P_\rho$ or $P_{f\rho}$, using the same Fourier–Bessel basis, excluding the constant mode, and the same cutoff length scale $\lambda_c$ as for the full dataset.

This procedure was repeated for several values of $N$, and several random subsets were generated for each value of $N$. We then computed the mean and standard deviation of the pattern measure at fixed $N$. The resulting dependence on cell number was fitted using

$$P(N) = 1 + qN,$$

where $q = q_\rho$ for the density pattern measure and $q = q_{f\rho}$ for the apparent expression pattern measure. The fitted value of $q$ therefore provides an empirical estimate of the strength of the large-scale signal captured by the selected modes.

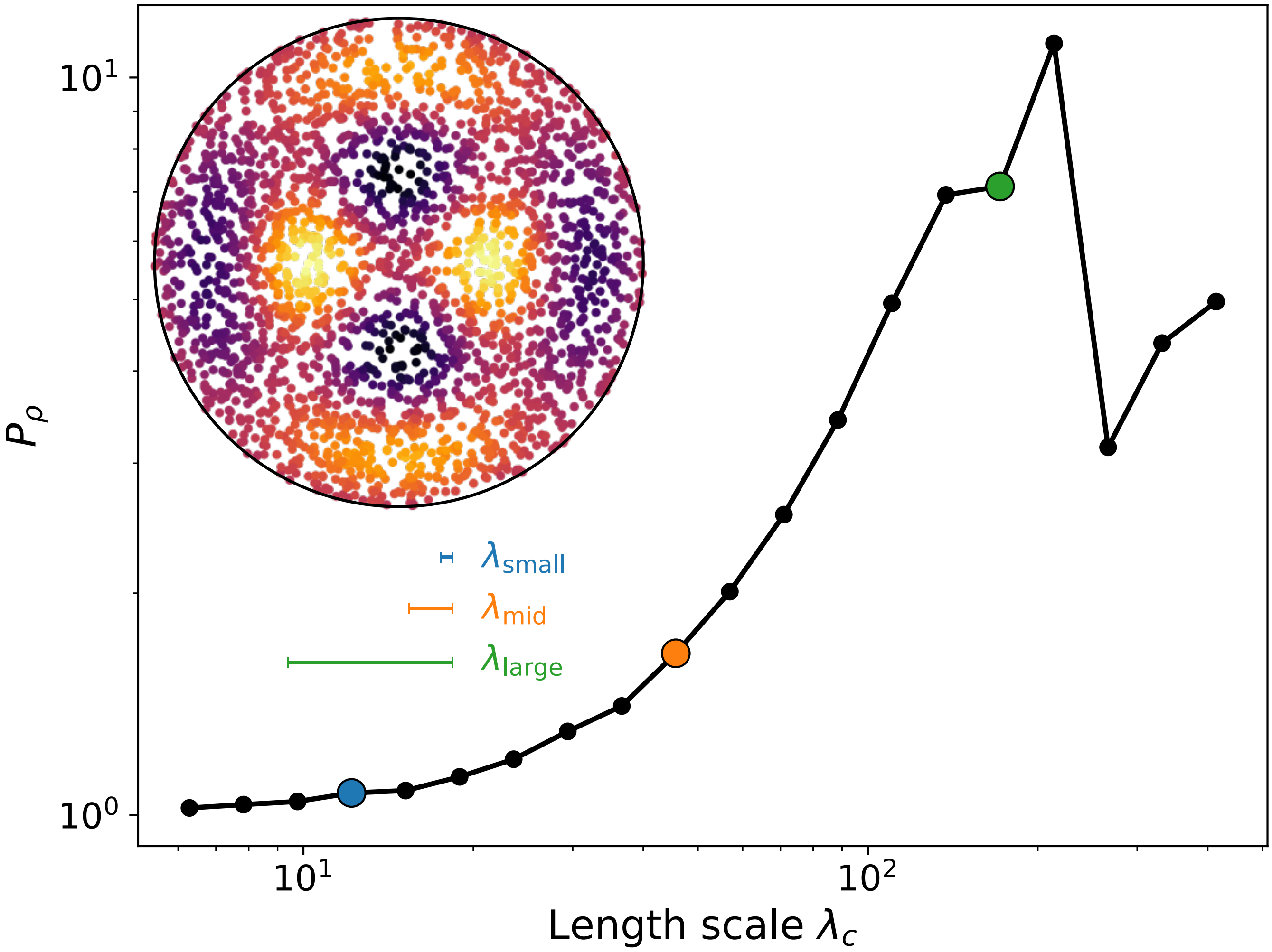

Figure S3: Dependence of the apparent expression pattern measure $P_{f\rho}$ on the cutoff length scale $\lambda_c$. Colored points mark three representative cutoff values; the matching bars in the inset show the corresponding length scales. The measure exhibits a pronounced maximum at an intermediate value of $\lambda_c$, indicating that the apparent expression pattern is dominated by a characteristic mode at that scale. For smaller $\lambda_c$, additional shorter-wavelength modes are included and reduce the average spectral contribution per retained mode.

#### 4 Dependence on the cutoff length scale

The pattern measures $P_\rho$ and $P_{f\rho}$ depend on the cutoff length scale $\lambda_c$, which defines which nonconstant Fourier–Bessel modes are included in the measure. Decreasing $\lambda_c$ includes progressively shorter-wavelength modes. Since the measures quantify the average spectral energy per retained mode, including additional short-wavelength modes can decrease the value of the measure when these modes contribute less strongly to the large-scale pattern.

For the density pattern measure $P_\rho$, this dependence is shown in the main text. The analogous analysis for the apparent expression pattern measure $P_{f\rho}$ is shown in Fig. S3. In this example, $P_{f\rho}$ displays a pronounced maximum at an intermediate cutoff length scale, indicating that the apparent expression pattern is dominated by modes at this characteristic scale. For smaller values of $\lambda_c$, additional shorter-wavelength modes are included, which reduces the average spectral energy per retained mode.

### 5 Exact temporal decomposition of spectral pattern measures

We consider the spectral pattern measure introduced in the main text, defined as

$$P_{f\rho}(t;\mu) = \frac{P_{\text{num}}(t;\mu)}{P_{\text{norm}}(t;\mu)}, \tag{1}$$

where $\mu \in \{0,1\}$ specifies whether the scalar cell-associated signal is centered with respect to its instantaneous population mean. For $\mu = 0$, the uncentered signal $f_k(t)$ is used, whereas for $\mu = 1$, the centered signal $f_k(t) - \langle f\rangle(t)$ is considered.

The numerator is given by

$$P_{\text{num}}(t;\mu) = \sum_{\substack{z\in\mathcal{Z}^* \\ \lambda_z>\lambda_c}} \left| d_z^{(\mu)}(t)\right|^2, \tag{2}$$

and the normalization term is defined as

$$P_{\text{norm}}(t;\mu) = \frac{M_c}{\pi R^2} \sum_{k\in\Omega(t)} \left(f_k(t) - \mu\langle f\rangle(t)\right)^2, \tag{3}$$

where $M_c = \#\{z \in \mathcal{Z}^* : \lambda_z > \lambda_c\}$ denotes the number of nonconstant Fourier–Bessel modes included in the numerator.

For each discrete time point $t_i$, the Fourier–Bessel coefficients are written in the form

$$d_z^{(\mu)}(t_i) := d_z^i = \frac{1}{\sqrt{K_z}} S_\mu^i, \quad z = (m,n) \in \mathcal{Z}^*, \qquad S_\mu^i := \sum_{k\in\Omega(t_i)} \left(f_k^i - \mu\langle f\rangle^i\right) g_z(r_k^i)\, e^{-im\theta_k^i}, \tag{4}$$

where $\Omega(t_i)$ denotes the set of tracked cells at time $t_i$, and $g_z(r) := J_{|m|}\big(\beta_{|m|,n}\, r/R\big)$ corresponds to the radial part of the Fourier–Bessel basis. The quantity $K_z$ denotes the normalization constant associated with the corresponding Fourier–Bessel mode.

The quantity $f_k^i$ denotes the scalar value associated with cell $k$ at time $t_i$, such as a gene expression level. The population average at time $t_i$ is defined as

$$\langle f\rangle^i := \frac{1}{|\Omega(t_i)|} \sum_{k\in\Omega(t_i)} f_k^i.$$

For later convenience, we introduce the notation $N^i := |\Omega(t_i)|$ for the number of tracked cells and $F^i := \sum_{k\in\Omega(t_i)} f_k^i$ for the corresponding total signal, so that $\langle f\rangle^i = F^i/N^i$.

#### 5.1 Time-symmetric decomposition of the numerator

##### Decomposition of $\langle f\rangle^{i+1} - \langle f\rangle^i$

We consider the discrete-time variation of the population average $\langle f\rangle^i = F^i/N^i$, which can be written exactly as

$$\langle f\rangle^{i+1} - \langle f\rangle^i = \frac{F^{i+1}}{N^{i+1}} - \frac{F^i}{N^i}. \tag{5}$$

Using a time-symmetric rearrangement, this difference can be expressed as

$$\langle f\rangle^{i+1} - \langle f\rangle^i = \frac{1}{2}\left(\frac{1}{N^{i+1}} + \frac{1}{N^i}\right)(F^{i+1} - F^i) + \frac{1}{2}\,\frac{F^{i+1}+F^i}{N^{i+1}N^i}\,(N^i - N^{i+1}), \tag{6}$$

which separates contributions associated with changes in the total signal and changes in the population size. This identity is exact and does not rely on any small-time approximation.

To interpret these terms, we partition the sets of tracked cells according to their presence at successive time points. Cells present at both times contribute to intrinsic variations of the scalar quantity, while appearing and disappearing cells contribute through changes in the total signal and population size. We further distinguish appearance and disappearance events associated with cell division from those not associated with division.

With this partition, the variation of the total signal can be written as

$$\begin{aligned} F^{i+1} - F^i = & \sum_{k\in\Omega(t_{i+1})\cap\Omega(t_i)} (f_k^{i+1} - f_k^i) \\ & + \left( \sum_{k\in\Omega_{\text{app}}(t_{i+1})} f_k^{i+1} - \sum_{k\in\Omega_{\text{disapp}}(t_i)} f_k^i \right) \\ & + \left( \sum_{k\in\Omega_{\text{daughters}}(t_{i+1})} f_k^{i+1} - \sum_{k\in\Omega_{\text{mother}}(t_i)} f_k^i \right). \end{aligned} \tag{7}$$

and the corresponding variation of the population size reads

$$N^{i+1} - N^i = \left|\Omega_{\text{app}}(t_{i+1})\right| - \left|\Omega_{\text{disapp}}(t_i)\right| + \left|\Omega_{\text{daughters}}(t_{i+1})\right| - \left|\Omega_{\text{mother}}(t_i)\right|. \tag{8}$$

Substituting these expressions into Eq. (6) yields an exact additive decomposition of the population-average variation,

$$\langle f\rangle^{i+1} - \langle f\rangle^i = E^{\langle\cdot\rangle} + D^{\langle\cdot\rangle} + B^{\langle\cdot\rangle}, \tag{9}$$

where $E^{\langle\cdot\rangle}$ quantifies the contribution of intrinsic changes in the cell-associated scalar quantity (e.g. gene expression levels) for cells tracked at both time points, $D^{\langle\cdot\rangle}$ captures contributions associated with cell division events, and $B^{\langle\cdot\rangle}$ accounts for contributions arising from cell appearance or disappearance not associated with division.

##### Decomposition of $S_\mu^{i+1} - S_\mu^i$ and of $d_z^{i+1} - d_z^i$

We consider the discrete-time increment of the quantity $S_\mu^i$ defined in the previous section. As for the decomposition of the population average, we partition the set of cells according to their presence at successive time points, separating contributions from cells tracked at both times, from those associated with cell appearance or disappearance.

Accordingly, the increment $S_\mu^{i+1} - S_\mu^i$ can be written as the sum of three terms, corresponding respectively to cells present at both time points, to appearing cells, and to disappearing cells. We focus first on the contribution of cells tracked at both times.

For each tracked cell $k$, we introduce the notation

$$M_k^j := g_z(r_k^j)\, e^{-im\theta_k^j}, \qquad \tilde{f}_{\mu,k}^j := f_k^j - \mu\langle f\rangle^j, \tag{10}$$

so that the contribution of tracked cells reads

$$\sum_{k\in\Omega(t_{i+1})\cap\Omega(t_i)} \left( \tilde{f}_{\mu,k}^{i+1} M_k^{i+1} - \tilde{f}_{\mu,k}^i M_k^i \right). \tag{11}$$

For each tracked cell, this difference is rewritten using the exact midpoint identity, yielding the time-symmetric expression

$$\tilde{f}_{\mu,k}^{i+1} M_k^{i+1} - \tilde{f}_{\mu,k}^i M_k^i = (\tilde{f}_{\mu,k}^{i+1} - \tilde{f}_{\mu,k}^i)\,\frac{M_k^{i+1} + M_k^i}{2} + (M_k^{i+1} - M_k^i)\,\frac{\tilde{f}_{\mu,k}^{i+1} + \tilde{f}_{\mu,k}^i}{2}. \tag{12}$$

Expanding $\tilde{f}^j_{\mu,k} = f^j_k - \mu\langle f\rangle^j$ and using the decomposition of $\langle f\rangle^{i+1} - \langle f\rangle^i$ derived above, the full increment $S^{i+1}_\mu - S^i_\mu$ admits an exact additive decomposition of the form

$$S^{i+1}_\mu - S^i_\mu = E^{(S)}_\mu + M^{(S)}_\mu + D^{(S)}_\mu + B^{(S)}_\mu. \tag{13}$$

Here, $E^{(S)}_\mu$ quantifies the contribution of intrinsic changes in the cell-associated scalar quantity (e.g. gene expression levels) for cells tracked at both time points, $M^{(S)}_\mu$ captures the contribution arising from cell motility through positional changes, $D^{(S)}_\mu$ accounts for contributions associated with cell division events, and $B^{(S)}_\mu$ accounts for contributions due to cell appearance or disappearance not associated with division.

Finally, since the spectral coefficients satisfy $d^i_z = S^i_\mu/\sqrt{K_z}$, the same decomposition applies to their temporal increments,

$$d^{i+1}_z - d^i_z = \frac{E^{(S)}_\mu + M^{(S)}_\mu + D^{(S)}_\mu + B^{(S)}_\mu}{\sqrt{K_z}}. \tag{14}$$

**Conclusion**

Collecting the previous results, the discrete-time increment of the numerator can be written as

$$P^{i+1}_{\text{num}} - P^i_{\text{num}} = \sum_{\substack{z\in\mathcal{Z}^* \\ \lambda_z>\lambda_c}} \Re\Big[(d^{i+1}_z - d^i_z)(\overline{d^{i+1}_z} + \overline{d^i_z})\Big]. \tag{15}$$

#### 5.2 Time-symmetric decomposition of the denominator

We consider the discrete-time increment of the normalization term

$$P^i_{\text{norm}} = \frac{M_c}{\pi R^2} \sum_{k\in\Omega(t_i)} (\tilde{f}^i_{\mu,k})^2, \qquad \tilde{f}^i_{\mu,k} := f^i_k - \mu\langle f\rangle^i.$$

Its discrete-time variation reads

$$P^{i+1}_{\text{norm}} - P^i_{\text{norm}} = \frac{M_c}{\pi R^2}\left[\sum_{k\in\Omega(t_{i+1})} (\tilde{f}^{i+1}_{\mu,k})^2 - \sum_{k\in\Omega(t_i)} (\tilde{f}^i_{\mu,k})^2\right].$$

Using the same partition of the cell sets as introduced previously, we separate contributions from cells tracked at both time points, from appearance, disappearance, and division events. This leads to the exact decomposition

$$P^{i+1}_{\text{norm}} - P^i_{\text{norm}} = E^{\text{norm}}_\mu + D^{\text{norm}}_\mu + B^{\text{norm}}_\mu.$$

For cells tracked at both time points we use the identity

$$(\tilde{f}^{i+1}_{\mu,k})^2 - (\tilde{f}^i_{\mu,k})^2 = (\tilde{f}^{i+1}_{\mu,k} - \tilde{f}^i_{\mu,k})(\tilde{f}^{i+1}_{\mu,k} + \tilde{f}^i_{\mu,k}),$$

together with

$$\tilde{f}^{i+1}_{\mu,k} - \tilde{f}^i_{\mu,k} = (f^{i+1}_k - f^i_k) - \mu(\langle f\rangle^{i+1} - \langle f\rangle^i).$$

Using the decomposition of the population average derived previously,

$$\langle f\rangle^{i+1} - \langle f\rangle^i = E^{\langle\cdot\rangle} + D^{\langle\cdot\rangle} + B^{\langle\cdot\rangle},$$

the normalization increment can be written as the sum of intrinsic expression changes, division contributions, and appearance/disappearance contributions.

#### 5.3 Time-symmetric decomposition of $P_{f\rho}^{i+1} - P_{f\rho}^{i}$

We now derive the exact discrete-time variation of the pattern measure

$$P_{f\rho}^{i} = \frac{P_{\text{num}}^{i}}{P_{\text{norm}}^{i}}.$$

Using a midpoint (time-symmetric) quotient identity, the discrete-time variation can be written exactly as

$$P_{f\rho}^{i+1} - P_{f\rho}^{i} = \frac{\overline{P_{\text{norm}}}\left(P_{\text{num}}^{i+1} - P_{\text{num}}^{i}\right) - \overline{P_{\text{num}}}\left(P_{\text{norm}}^{i+1} - P_{\text{norm}}^{i}\right)}{P_{\text{norm}}^{i+1} P_{\text{norm}}^{i}}, \qquad \overline{X} := \frac{X^{i+1} + X^{i}}{2}. \tag{16}$$

This identity is exact and does not rely on any small-time approximation.

The increments $P_{\text{num}}^{i+1} - P_{\text{num}}^{i}$ and $P_{\text{norm}}^{i+1} - P_{\text{norm}}^{i}$ were derived in the previous sections. Substituting these expressions yields the temporal decomposition

$$P_{f\rho}(t_{i+1};\mu) - P_{f\rho}(t_i;\mu) = E^{i+\frac{1}{2}} + M^{i+\frac{1}{2}} + D^{i+\frac{1}{2}} + B^{i+\frac{1}{2}}.$$

Here, the terms $E$, $M$, $D$, and $B$ correspond respectively to the contributions of intrinsic changes in the cell-associated scalar quantity, cell motility, cell division, and appearance or disappearance events not associated with division.

These decompositions are exact at finite time steps and provide a direct link between the temporal evolution of the pattern measure and the underlying cellular processes.

#### 5.4 Temporal decomposition of the pattern measure

From the previous sections, the discrete-time variation of the pattern measure satisfies

$$P_{f\rho}^{i+1} - P_{f\rho}^{i} = E^{i+\frac{1}{2}} + M^{i+\frac{1}{2}} + D^{i+\frac{1}{2}} + B^{i+\frac{1}{2}}.$$

Since this identity holds for every experimental time interval $[t_i, t_{i+1}]$, the variation of the pattern measure between the initial time $t_0$ and any later time $t$ can be obtained by summing the contributions over successive time steps. Using the telescopic identity

$$P_{f\rho}(t) - P_{f\rho}(t_0) = \sum_{i:\, t_i \in [t_0, t)} \left(P_{f\rho}^{i+1} - P_{f\rho}^{i}\right),$$

we obtain

$$P_{f\rho}(t) - P_{f\rho}(t_0) = \sum_{i:\, t_i \in [t_0, t)} \left(E^{i+\frac{1}{2}} + M^{i+\frac{1}{2}} + D^{i+\frac{1}{2}} + B^{i+\frac{1}{2}}\right).$$

We then define the cumulative contribution associated with each cellular process $X \in \{E, M, D, B\}$ as

$$\mathcal{X}(t) := \sum_{i:\, t_i \in [t_0, t)} X^{i+\frac{1}{2}},$$

so that the total variation of the pattern measure can be written as

$$P_{f\rho}(t) - P_{f\rho}(t_0) = \mathcal{E}(t) + \mathcal{M}(t) + \mathcal{D}(t) + \mathcal{B}(t).$$

This expression provides an exact cumulative decomposition of the change in the pattern measure from the initial configuration into contributions arising from expression dynamics, cell motility, cell division, and cell appearance or disappearance.

For each cellular process $X \in \{E, M, D, B\}$, the relative contribution to the net change in the pattern measure at the final observation time $t_f$ is defined as

$$R_X = 100 \times \frac{\mathcal{X}(t_f)}{P_{f\rho}(t_f) - P_{f\rho}(t_0)}.$$

##### 5.5 Refined decomposition when boundary-crossing events can be identified

In the decomposition described above, the term $B$ accounts for cell appearance and disappearance events that are not associated with cell division. When the data contain sufficient information to determine whether cells enter or leave the analyzed region through its boundary, this contribution can be further decomposed.

This occurs, for instance, when tracking is performed on a larger field of view than the circular region used for the Fourier–Bessel decomposition. In our case, the original data are defined on a square domain, while the spectral analysis is restricted to a circular region of radius $R$. This makes it possible to distinguish true appearance and disappearance events from boundary-crossing events.

We therefore further decompose the sets of non-division-related appearance and disappearance events as

$$\Omega_{\text{app}}(t_{i+1}) = \Omega_{\text{app}}^{\text{true}}(t_{i+1}) \cup \Omega_{\text{in}}^{\partial\Omega}(t_{i+1}),$$

and

$$\Omega_{\text{disapp}}(t_i) = \Omega_{\text{disapp}}^{\text{true}}(t_i) \cup \Omega_{\text{out}}^{\partial\Omega}(t_i),$$

where $\Omega_{\text{in}}^{\partial\Omega}$ denotes cells entering the circular region through its boundary, and $\Omega_{\text{out}}^{\partial\Omega}$ denotes cells leaving the circular region through its boundary.

Accordingly, the balance contribution $B$ can be further decomposed into two terms,

$$B = B_{\text{app/disapp}} + B_{\partial\Omega},$$

where $B_{\text{app/disapp}}$ accounts for true appearance and disappearance events, while $B_{\partial\Omega}$ accounts for boundary-crossing events associated with cells entering or leaving the circular region of interest.

The rest of the decomposition is unchanged. In particular, the same splitting can be applied to the increments of the population average, the Fourier–Bessel coefficients, the numerator, the normalization factor, and therefore to the final pattern measure. This yields

$$P_{f\rho}^{i+1} - P_{f\rho}^{i} = E^{i+\frac{1}{2}} + M^{i+\frac{1}{2}} + D^{i+\frac{1}{2}} + B_{\text{app/disapp}}^{i+\frac{1}{2}} + B_{\partial\Omega}^{i+\frac{1}{2}},$$

where $B_{\partial\Omega}^{i+\frac{1}{2}}$ isolates the contribution due to boundary fluxes across the circular region of interest.

### 6 Agent-based model details

This section provides the full technical description of the agent-based models (ABMs) used to validate the framework. These models are designed as controlled, phenomenological systems in which the mechanisms responsible for spatial pattern formation are explicitly prescribed. They are not intended to provide a detailed biophysical description of real biological systems, but rather to generate ground-truth scenarios for method validation.

We describe here two classes of ABMs. The first model considers agents characterized solely by their positions and allows spatial pattern to emerge from motility, division, and apoptosis. The second model extends this framework by coupling spatial dynamics to gene expression carried by individual agents. All simulations are performed in two spatial dimensions within a circular observation domain.

#### 6.1 Model without gene expression

**Agent positions and domain geometry** Agents are represented as point-like particles evolving in a two-dimensional continuous space. The simulation domain is a disk of radius $R = L/2$, centered at the origin. Let $\mathbf{x}_i(t) \in \mathbb{R}^2$ denote the position of agent $i$ at time $t$.

**Equations of motion** The total deterministic force $\mathbf{F}_i$ acting on agent $i$ includes contributions from pairwise interactions with neighboring agents and from external confining forces. Agent positions evolve according to overdamped Langevin dynamics. The temporal evolution of each agent position is given by

$$\mathbf{x}_i(t + \Delta t) = \mathbf{x}_i(t) + \mu \mathbf{F}_i(t)\Delta t + \sqrt{2D_x \Delta t}\, \boldsymbol{\xi}_i(t),$$

where $\mu$ is the mobility, $D_x$ is the translational diffusion coefficient, and $\boldsymbol{\xi}_i(t)$ is a two-dimensional vector of independent Gaussian random variables with zero mean and unit variance.

**Confining boundary force** To ensure spatial confinement within the circular domain while avoiding hard boundary conditions, agents experience a soft radial restoring force when approaching the domain boundary. Let $r_i = \|\mathbf{x}_i\|$ denote the radial position of agent $i$. When $r_i$ exceeds a threshold radius $r_0 < R$, a confining force is applied according to

$$\mathbf{F}_i^{\text{wall}} = -2k_{\text{wall}}(r_i - r_0)\,\frac{\mathbf{x}_i}{r_i},$$

where $k_{\text{wall}}$ sets the stiffness of the confinement. No boundary force is applied for $r_i \leq r_0$. This soft confinement biases agent motion toward the interior of the domain while preserving smooth dynamics near the boundary.

**Pairwise mechanical interactions** Mechanical interactions between agents consist of a short-range repulsive component and a longer-range adhesive component. These interactions provide a minimal and controlled implementation of spatial pattern formation that is due to motility only.

For a pair of agents $i$ and $j$ separated by a distance $r_{ij} = \|\mathbf{x}_i - \mathbf{x}_j\|$, short-range repulsion is modeled as the gradient of a Gaussian interaction kernel. The repulsive force exerted on agent $i$ by agent $j$ is given by

$$\mathbf{F}_{ij}^{\text{rep}} = \varepsilon\,\frac{1}{\sigma_N^2}\frac{1}{2\pi\sigma_N^2}\exp\left(-\frac{r_{ij}^2}{2\sigma_N^2}\right)(\mathbf{x}_i - \mathbf{x}_j),$$

and is applied only for $r_{ij} \leq r_{\text{rep}}$, where $r_{\text{rep}}$ is a cutoff radius proportional to $\sigma_N$. The interaction length scale $\sigma_N$ depends weakly on the total number of agents $N$ according to

$$\sigma_N = \ell_{r0} N^{-\beta/2},$$

which allows the effective interaction range to adapt to changes in population size.

Long-range adhesion is modeled as an attractive interaction with an exponentially decaying amplitude. The adhesive force exerted on agent $i$ by agent $j$ is defined as

$$\mathbf{F}_{ij}^{\mathrm{adh}} = -\frac{A}{\ell_a} \exp\left(-\frac{r_{ij}}{\ell_a}\right) \frac{\mathbf{x}_i - \mathbf{x}_j}{r_{ij}},$$

and is applied for $r_{ij} \leq r_{\mathrm{adh}}$, where $r_{\mathrm{adh}}$ denotes the adhesion cutoff radius. To avoid discontinuities in the force, the adhesive interaction is smoothly tapered to zero near $r_{\mathrm{adh}}$.

**Mechanical mixing parameter** The strengths of repulsive and adhesive interactions are controlled by a dimensionless mechanical mixing parameter $w \in [0, 1]$. Both interaction amplitudes $\varepsilon$ and $A$ are rescaled by a common factor proportional to $(1 - w)^\eta$, with $\eta > 1$. Mechanical interactions act at full strength for $w = 0$ and are progressively suppressed as $w$ approaches one, leaving only stochastic motion, confinement, and population turnover.

**Local density estimation** Structured population turnover relies on an estimate of the local agent density. For each agent $i$, a local density $\rho_i$ is computed as a weighted sum over neighboring agents within a finite interaction radius. Specifically,

$$\rho_i = \sum_{j \neq i} K(\|\mathbf{x}_j - \mathbf{x}_i\|),$$

where the kernel $K(r)$ is either uniform within a cutoff radius or Gaussian, depending on the process considered. To account for boundary effects, density estimates are renormalized by the fraction of the kernel support that lies within the observation domain. This correction prevents systematic underestimation of local density near the boundary.

**Division dynamics** Agent division is implemented as a Poisson process with an instantaneous rate that depends on short-range local density. For each agent $i$, we compute an effective local density $\rho_i$ within a radius $R_{\mathrm{div}}$ using the kernel-based estimator described above. The structured division rate is defined as

$$\lambda_i^{\mathrm{div,struct}} = \lambda_{\mathrm{div}}\, f\left(\frac{\rho_i}{K_{\mathrm{div}}}\right), \qquad f(x) = \frac{x^h}{1 + x^h},$$

where $\lambda_{\mathrm{div}}$ sets the maximal structured division rate, $K_{\mathrm{div}}$ is a characteristic density scale, and $h \geq 1$ is a Hill exponent.

Division is then blended with a spatially uniform baseline turnover rate using the turnover mixing parameter $\tau \in [0, 1]$,

$$\lambda_i^{\mathrm{div}} = (1 - \tau)\lambda^{\mathrm{div,rand}} + \tau \lambda_i^{\mathrm{div,struct}}.$$

This construction ensures that a spatially uniform baseline division rate is always present, while additional density-dependent structure emerges once a sufficiently populated local neighborhood is present.

When a division event occurs, the parent agent is removed and replaced by two daughter agents placed near the parent location with small Gaussian displacements. Parent–daughter relationships are recorded through persistent identifiers in the exported trajectories.

**Apoptosis dynamics** Apoptosis is implemented as a stochastic death process whose rate depends on local density evaluated over a longer spatial range. For each agent $i$, an effective long-range density $\rho_i$ is computed using the kernel-based estimator described above. The structured apoptosis rate is defined as

$$\lambda_i^{\mathrm{death,struct}} = \lambda_0^{\mathrm{death}} + \lambda_{\mathrm{crowd}}^{\mathrm{death}}\, g\left(\frac{\rho_i}{K_{\mathrm{death}}}\right), \qquad g(y) = y^p,$$

where $\lambda_0^{\mathrm{death}}$ is a constant offset, $\lambda_{\mathrm{crowd}}^{\mathrm{death}}$ controls the strength of density-dependent death, $K_{\mathrm{death}}$ is a reference density scale, and $p$ is a positive exponent controlling the sensitivity of apoptosis to crowding.

As for division, apoptosis rates are blended with a spatially uniform background rate using the turnover mixing parameter $\tau$,

$$\lambda_i^{\mathrm{death}} = (1-\tau)\lambda^{\mathrm{death,rand}} + \tau\lambda_i^{\mathrm{death,struct}}.$$

This formulation introduces long-range, density-dependent apoptosis while maintaining a uniform background turnover.

**Turnover mixing parameter** The relative contribution of structured population turnover is controlled by a turnover mixing parameter $\tau \in [0, 1]$. For both division and apoptosis, instantaneous rates are defined as a convex combination of a uniform, spatially unstructured rate and the structured, density-dependent rate described above. Specifically,

$$\lambda_i = (1-\tau)\lambda^{\mathrm{rand}} + \tau\lambda_i^{\mathrm{struct}}.$$

For $\tau = 0$, division and apoptosis occur uniformly in space and do not contribute to spatial pattern. For $\tau = 1$, turnover is fully structured and can generate spatial patterns independently of motility. Intermediate values correspond to mixed regimes in which turnover and motility jointly contribute to pattern formation.

**Relation to the mixing parameter $\alpha$** For clarity, spatial pattern formation is described in the manuscript using a single mixing parameter $\alpha \in [0, 1]$. In the agent-based models, this parameter corresponds to a specific one-dimensional slice of the internal parameter space defined by the mechanical mixing parameter $w$ and the turnover mixing parameter $\tau$.

For the validation presented here, we restrict to the diagonal choice

$$w = \tau = \alpha,$$

which interpolates continuously between a purely motility-driven regime ($\alpha = 0$) and a purely turnover-driven regime ($\alpha = 1$). This construction is illustrated schematically in Fig. S4. Intermediate values of $\alpha$ correspond to mixed regimes in which both motility and turnover contribute to spatial pattern. More general combinations of $w$ and $\tau$ are possible but are not detailed here.

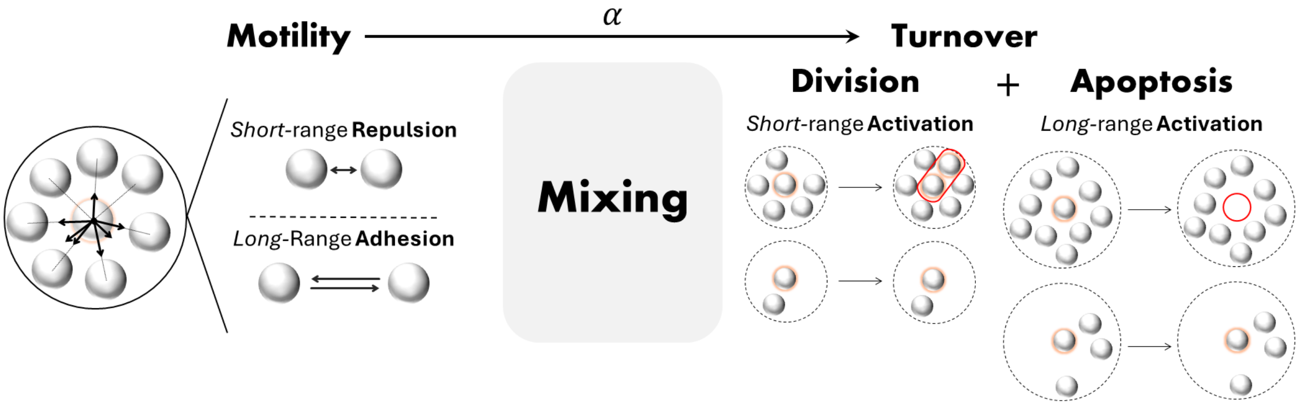

Figure S4: Schematic of the agent-based model, combining cell motility and structured population turnover through division and apoptosis, with their relative contribution controlled by a mixing parameter $\alpha$.

#### 6.2 Model with gene expression dynamics

**State variables and overview** In this second class of agent-based models, each agent carries both a position $\mathbf{x}_i(t) \in \mathbb{R}^2$ and a continuous internal state $s_i(t) \in \mathbb{R}$ representing gene expression. Introducing an internal gene expression state expands the space of possible mechanisms by which spatial patterns can emerge in these models. In the following, we focus on regimes in which patterning arises from the interplay between cell motility and expression-dependent interactions. In the simulations reported in this work, cell division and apoptosis are suppressed, such that spatial pattern arises solely from motility and expression. We verified in additional simulations not shown here that introducing rare division and apoptosis events does not qualitatively affect the results. The simulation domain is a disk of radius $R = L/2$, with soft confinement implemented through a radial wall force as described above.

**Expression dynamics** Each agent carries a continuous expression state $s_i(t) \in \mathbb{R}$. Expression dynamics are built upon an overdamped Langevin process in a symmetric double-well potential with minima at $\pm 1$, supplemented by additional interaction-induced drift terms.

$$U(s) = \frac{\alpha_{\rm dw}}{4}(s^2 - 1)^2, \qquad U'(s) = \alpha_{\rm dw}\, s(s^2 - 1).$$

Using an Euler–Maruyama discretization with time step $\Delta t$, expression is updated as

$$s_i(t + \Delta t) = s_i(t) + \Delta t \Big[-\kappa_s\, U'\big(s_i(t)\big) + \psi\, \eta_{\rm act}\big[\bar{s}_i^{\rm act}(t) - s_i(t)\big] - \psi\, \eta_{\rm inh}\big[\bar{s}_i^{\rm inh}(t) - s_i(t)\big]\Big] + \sqrt{2D_s\, \Delta t}\, \xi_i(t),$$

where $\eta_{\rm act}$ and $\eta_{\rm inh}$ are constant coupling coefficients that set the relative strengths of short-range activation and long-range inhibition, respectively, $\xi_i(t) \sim \mathcal{N}(0, 1)$ are independent Gaussian random variables, $D_s$ controls expression fluctuations, and $\psi \in [0, 1]$ sets the overall strength of neighborhood coupling. In the simulations reported here, $\psi$ is specified as a function of the mixing parameter $\alpha$.

The short-range and long-range neighborhood averages are computed as normalized, distance-weighted means,

$$\bar{s}_i^{\rm act} = \frac{\sum_{j\neq i} s_j\, K_{\rm act}(r_{ij})}{\sum_{j\neq i} K_{\rm act}(r_{ij})}, \qquad \bar{s}_i^{\rm inh} = \frac{\sum_{j\neq i} s_j\, K_{\rm inh}(r_{ij})}{\sum_{j\neq i} K_{\rm inh}(r_{ij})},$$

with $r_{ij} = \|\mathbf{x}_i - \mathbf{x}_j\|$, where $K_{\rm act}$ and $K_{\rm inh}$ are radially symmetric kernels with finite support and distinct spatial ranges. The two terms proportional to $\eta_{\rm act}$ and $\eta_{\rm inh}$ respectively promote short-range alignment of expression and counteract local homogenization over a longer range. To avoid rare numerical excursions induced by stochastic integration, expression values are bounded to a finite interval by truncation.

**Position dynamics with expression-dependent interactions** Agent positions evolve according to overdamped Langevin dynamics,

$$\mathbf{x}_i(t + \Delta t) = \mathbf{x}_i(t) + \mu \mathbf{F}_i(t)\Delta t + \sqrt{2D_x \Delta t}\, \boldsymbol{\xi}_i(t),$$

where $\mu$ is the mobility, $D_x$ the positional diffusion coefficient, and $\boldsymbol{\xi}_i(t)$ a two-dimensional Gaussian noise with zero mean and unit variance. The total force $\mathbf{F}_i$ includes pairwise short-range exclusion, longer-range adhesion, and the confining boundary force.

**Expression-dependent modulation of mechanical interactions** Mechanical interactions retain the same functional form as in the position-only model, consisting of short-range exclusion and longer-range adhesion. In the presence of gene expression, only the amplitudes of these interactions are modulated, while their spatial ranges and functional forms remain unchanged.

For each interacting pair $(i, j)$, we define the mean expression $m_{ij} = (s_i + s_j)/2$. Adhesion and exclusion amplitudes are then determined from a normalized weight $w(m_{ij}) \in [0, 1]$, constructed from an auxiliary asymmetric double-well mapping. This mapping provides a controlled phenomenological coupling between expression similarity and mechanical interactions, without introducing direct feedback on expression dynamics. To parameterize the expression dependence of interaction amplitudes, we introduce an asymmetric auxiliary energy function $f(m)$ defined for $m \in [-1, 1]$ as

$$f(m) = \frac{m^6}{6} - \frac{3\delta}{4}m^5 + \Big(h - \frac{1}{3}\Big)m^4 + \frac{5\delta}{4}m^3 - \Big(2h - \frac{1}{6}\Big)m^2,$$

where $h$ controls barrier height and $\delta$ controls asymmetry. To avoid hard clipping artifacts when $m_{ij}$ fluctuates outside the interval, we apply a smooth saturation $m_{\text{eff}} = \tanh(\gamma m_{ij})$ prior to evaluating $f$.

A normalized interaction weight is then defined as

$$w(m) = \text{clip}\left(1 - \frac{f(m_{\text{eff}}) - f_{\min}}{f_0 - f_{\min}}, \, 0, \, 1\right),$$

where $f_{\min} = \min\{f(-1), f(+1)\}$ and $f_0 = f(0)$. The clipping operation $\text{clip}(x, 0, 1) = \min(1, \max(0, x))$ enforces that interaction weights remain bounded within the interval $[0, 1]$.

Pairwise interaction amplitudes are then defined as

$$A_{ij} = A_{\min} + (A_{\max} - A_{\min})\, w(m_{ij})^{p_A}, \qquad \varepsilon_{ij} = \varepsilon_{\min} + (\varepsilon_{\max} - \varepsilon_{\min})\big(1 - w(m_{ij})\big)^{p_\varepsilon}.$$

Here, $A_{ij}$ denotes the amplitude of the adhesive force entering the long-range exponential attraction term, while $\varepsilon_{ij}$ controls the strength of the short-range repulsive interaction defined by the Gaussian kernel. This construction biases mechanical interactions based on expression similarity and can contribute to expression-dependent spatial segregation in combination with agent motion and stochastic fluctuations.

**Relation to the mixing parameter $\alpha$** In the simulations reported here, we use a single mixing parameter $\alpha \in [0, 1]$ to co-vary (i) the strength of neighborhood coupling in the expression dynamics and (ii) the degree of expression-dependence of mechanical interactions.

In the expression update rule defined above, the coupling to short-range activation and long-range inhibition terms is multiplied by

$$\psi(\alpha) = \alpha,$$

such that $\alpha = 0$ removes interaction-induced expression drift, yielding purely bistable expression dynamics with stochastic fluctuations, whereas $\alpha = 1$ yields maximal expression coupling.

For mechanical interactions, we interpolate between an expression-dependent interaction model and an expression-independent (constant-amplitude) model by defining, for each interacting pair $(i, j)$,

$$A_{ij}(\alpha) = (1 - \alpha)\, A_{ij}^{\text{expr}} + \alpha\, A_{\text{const}}, \qquad \varepsilon_{ij}(\alpha) = (1 - \alpha)\, \varepsilon_{ij}^{\text{expr}} + \alpha\, \varepsilon_{\text{const}},$$

where $A_{ij}^{\text{expr}}$ and $\varepsilon_{ij}^{\text{expr}}$ are the expression-dependent amplitudes defined above through the mapping $w(m_{ij})$, and $A_{\text{const}}, \varepsilon_{\text{const}}$ are constant amplitudes.

By construction, $\alpha = 0$ corresponds to a motility-driven regime in which gene expression evolves dynamically but does not contribute to spatial pattern, such that any emerging spatial structure arises from cell motion and mechanical sorting. In contrast, $\alpha = 1$ corresponds to a regime in which spatial pattern is driven by expression dynamics, while mechanical interactions are independent of expression. Intermediate values of $\alpha$ yield mixed regimes in which both mechanisms contribute to pattern formation.

##### 6.3 Numerical implementation

All simulations are performed using an explicit Euler–Maruyama discretization with a fixed time step $\Delta t$, using the same numerical scheme for both agent-based models, with and without gene expression dynamics.

At each time step, deterministic interaction forces are first computed from the current agent positions and, when applicable, their expression states. Agent positions are then updated according to the overdamped Langevin equation, including stochastic displacement. For models including gene expression, internal expression states are updated subsequently using the corresponding Langevin equation.

All simulations are initialized from spatially homogeneous configurations and run for a fixed number of time steps sufficient to reach a stationary or slowly evolving spatial pattering. Trajectories are recorded at every time step for downstream analysis.

### 7 Experimental data processing

#### 7.1 Presomitic mesoderm cell aggregates

We applied our framework to experimental single-cell tracking data obtained from two independent two-dimensional aggregates of mouse presomitic mesoderm cells [1]. These datasets consist of time-resolved trajectories of individual cells within approximately circular aggregates, obtained through live imaging and automated single-cell tracking.

###### Single-cell tracking data and preprocessing.

We analyzed the single-cell tracking datasets generated and characterized by Zhao et al. [2].

**Choice of the radial domain.** Presomitic mesoderm aggregates have an overall radial organization, with a pronounced density decrease near the aggregate boundary. If the analysis radius $R$ is chosen too large, the Fourier–Bessel measure can therefore capture this aggregate-scale boundary effect rather than spatial patterning within the aggregate.

To avoid this, we selected $R$ from the radial cell density profiles. For each time point, we computed the radially averaged density $\rho(r, t)$ and compared its evolution over time (Fig. S5). The profiles show a peripheral density drop beyond the aggregate interior. We therefore chose $R = 220$, which retains a large inner region while excluding the strongest boundary-associated density decrease. All subsequent Fourier–Bessel analyses of the presomitic mesoderm aggregates were performed on the disk $r < R$.

#### 7.2 Breast cancer cell populations

We applied the framework to single-cell tracking data obtained from live imaging experiments of breast cancer cell lines exhibiting spatial self-organization of cancer stem cell niches [3]. These data consist of time-resolved trajectories of individual cells obtained from long-term time-lapse microscopy of cultured breast cancer cell populations.

For the analysis presented here, we focused on two experiments performed on the SUM159PT breast cancer cell line. In the original dataset, cell trajectories are provided within a square observation domain corresponding to the microscope field of view. Because the spectral framework introduced in this work is defined on circular domains, we restricted the analysis to the largest circle fully contained within the observation window. Only cells located within this circular region were retained for the computation of the spectral pattern measure and its temporal decomposition. Unlike the presomitic mesoderm aggregates, these datasets do not exhibit an intrinsic aggregate-scale radial structure; the circular region was therefore used only to match the geometry of the Fourier–Bessel decomposition.

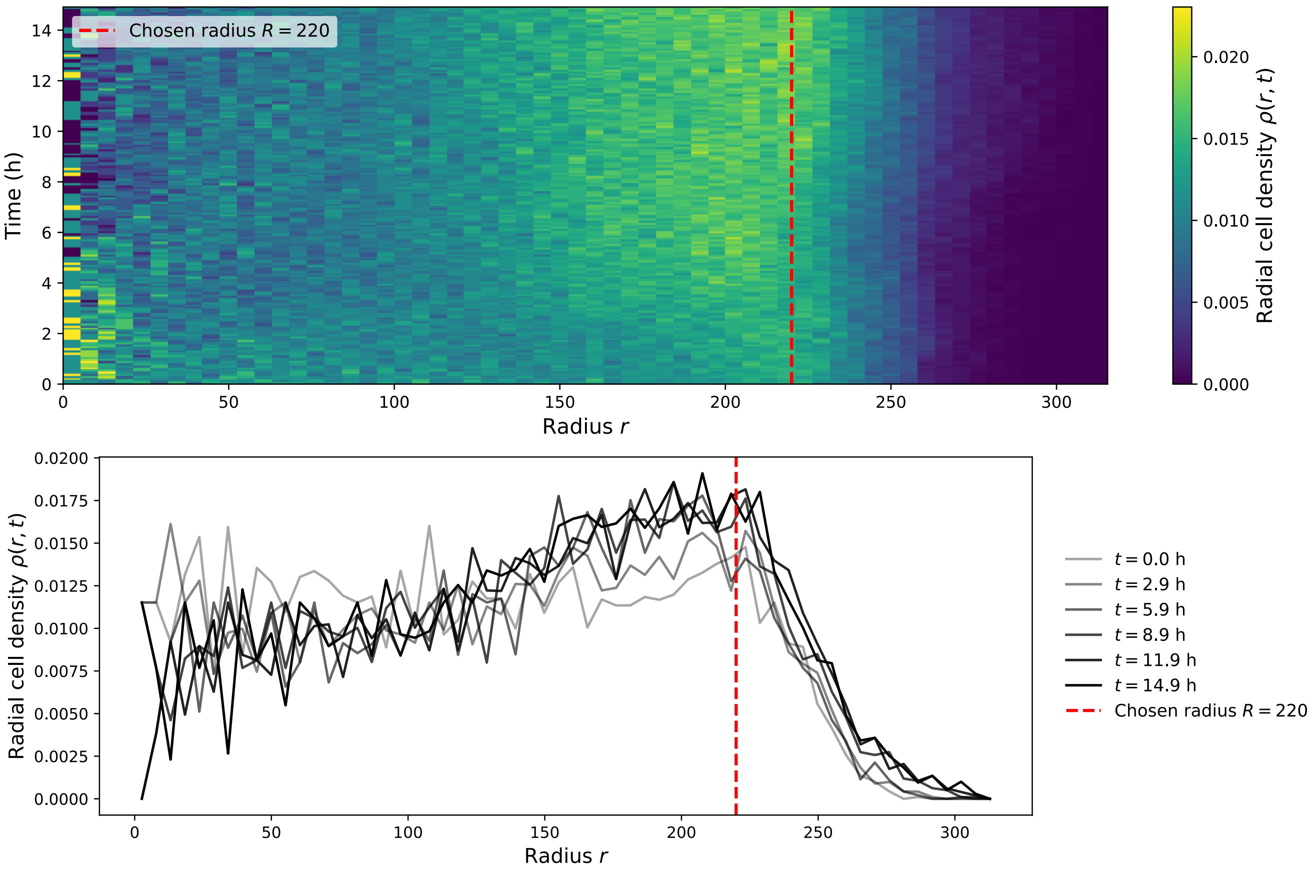

Figure S5: Choice of the radial domain for the analysis of presomitic mesoderm aggregates. (A) Radial cell density $\rho(r, t)$ as a function of radius and time. The dashed red line indicates the radius $R = 220$ used for the Fourier–Bessel analysis. (B) Radial density profiles at selected time points.

### 8 Sensitivity to the cutoff length scale

To assess whether the inferred cellular contributions depend on the choice of cutoff length scale, we repeated the temporal decomposition of Experiment 1 for several values of $\lambda_c$. For each value of $\lambda_c$, the cumulative contribution of each cellular process was evaluated at the final observation time $t_f$. The relative final contribution of each process was then obtained by normalizing its cumulative contribution at $t_f$ by the net change in the corresponding pattern measure, $P(t_f) - P(t_0)$. The dashed vertical line indicates the cutoff length scale used in the main analysis.

#### 8.1 Presomitic mesoderm aggregates (Exp. 1)

In the first presomitic mesoderm aggregate, the main qualitative conclusions remain robust over the tested range of cutoff length scales (Fig. S6). Around the selected value of $\lambda_c$, cell motility is the dominant positive contribution to both $P_\rho$ and $P_{f\rho}$. For $P_\rho$, boundary crossings also provide a substantial positive contribution, whereas dis-/appearance events contribute negatively. For $P_{f\rho}$, division provides an additional positive contribution at the final observation time, while the remaining contributions are comparatively smaller. Overall, the conclusion that pattern formation is primarily motility-driven is preserved.

#### 8.2 Breast cancer cell lines (Exp. 1)

In the first breast cancer experiment, the decomposition is also robust to the choice of $\lambda_c$, with cell division remaining the dominant positive contribution across the tested range (Fig. S7). For both $P_\rho$ and $P_{f\rho}$, dis-/appearance events provide an additional positive contribution, whereas boundary-crossing contributions remain close to zero. Motility contributes negatively, although its magnitude remains relatively small around the selected value of $\lambda_c$. For $P_{f\rho}$, expression changes contribute negatively over the tested range.

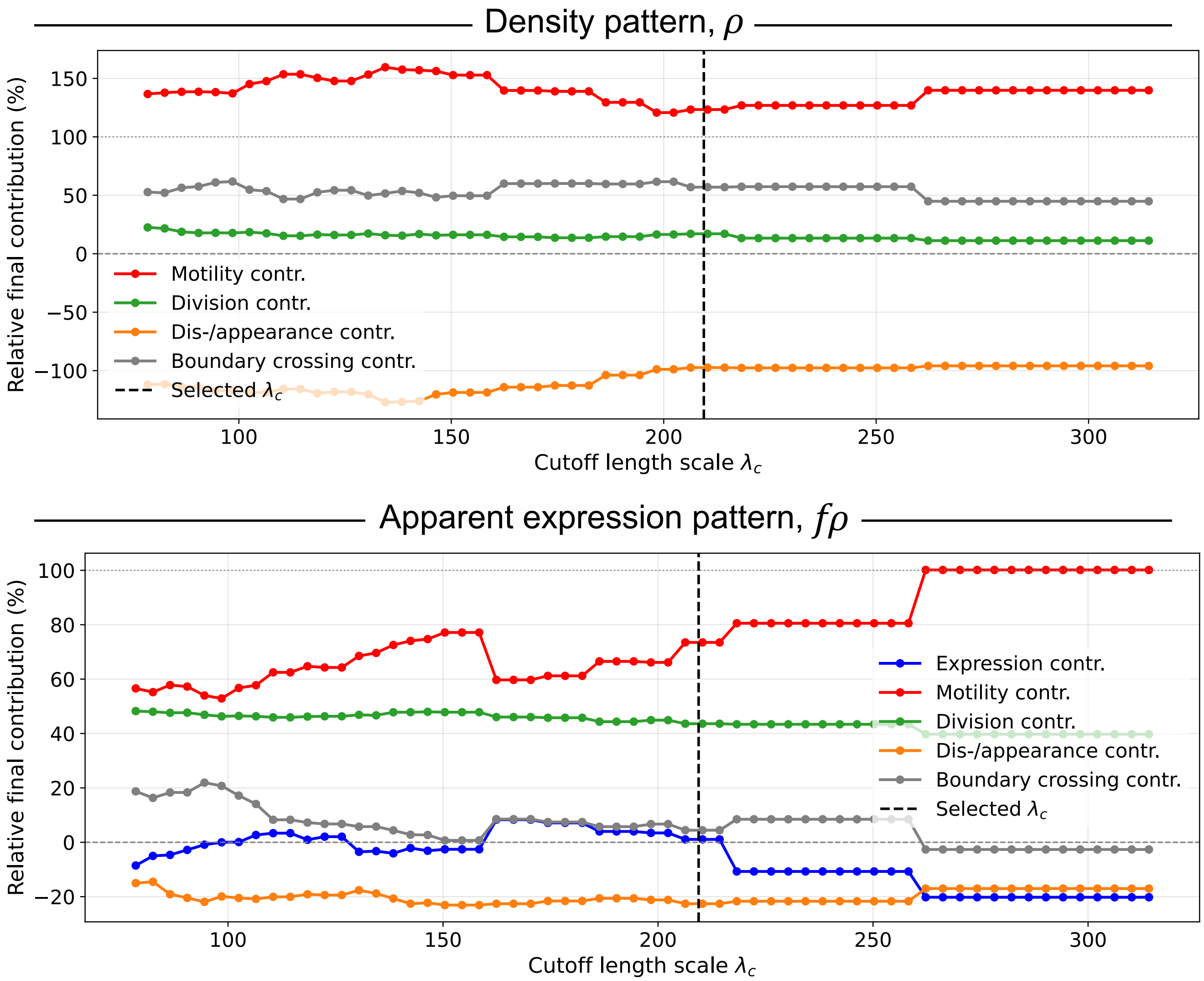

Figure S6: Sensitivity of the inferred cellular contributions to the cutoff length scale in the first presomitic mesoderm aggregate. Relative final contributions are shown as a function of $\lambda_c$ for the density pattern measure $P_\rho$ and the apparent expression pattern measure $P_{f\rho}$. The dashed vertical line indicates the cutoff length scale used in the main analysis. Across the tested range, cell motility remains the dominant positive contribution for both measures, while division, dis-/appearance, boundary crossing, and expression contributions remain secondary.

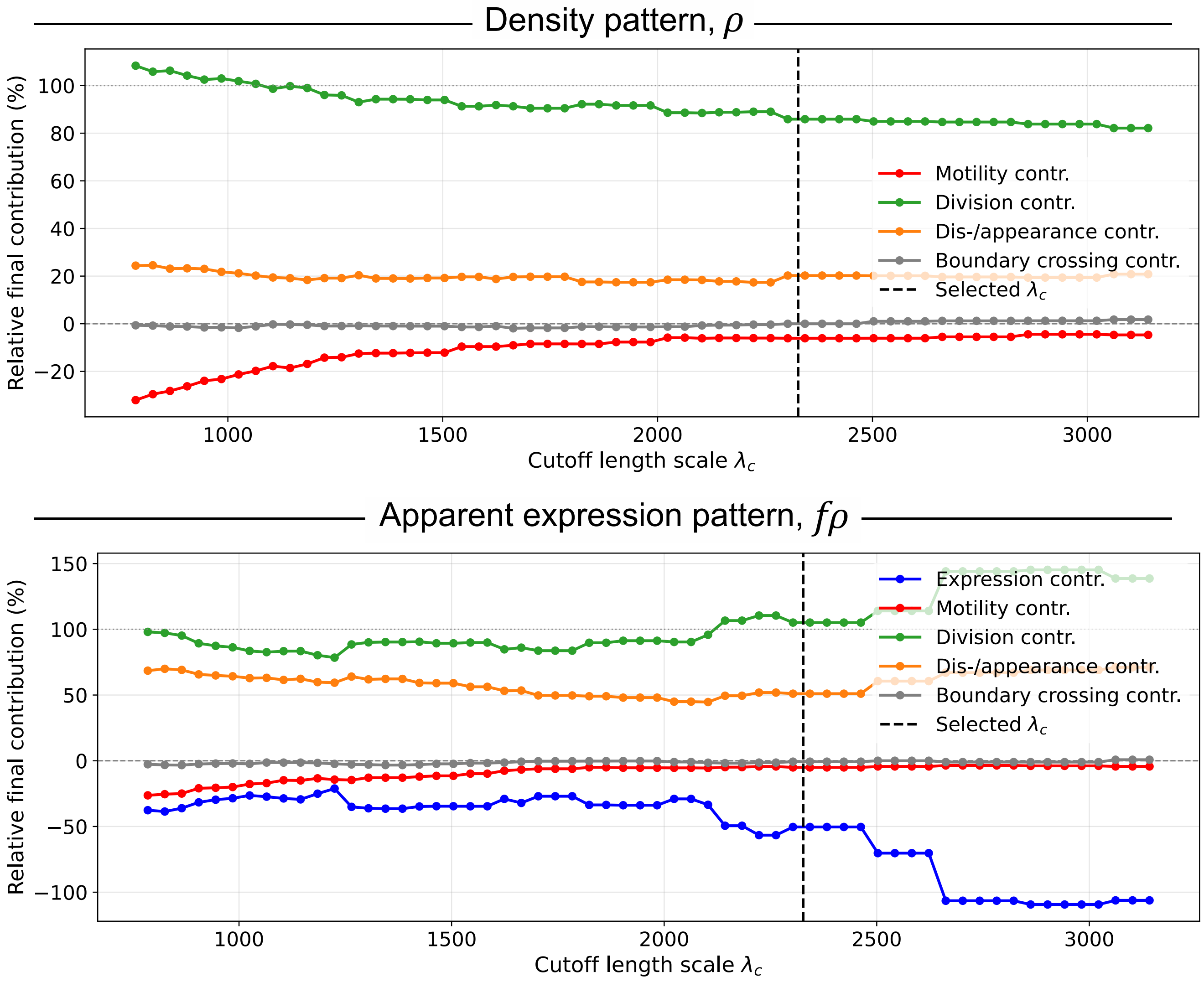

Figure S7: Sensitivity of the inferred cellular contributions to the cutoff length scale in the first breast cancer experiment. Relative final contributions are shown as a function of $\lambda_c$ for the density pattern measure $P_\rho$ and the apparent expression pattern measure $P_{f\rho}$. The dashed vertical line indicates the cutoff length scale used in the main analysis. Across the tested range, cell division remains the dominant positive contribution, while expression changes contribute negatively to $P_{f\rho}$. Motility and boundary crossing contributions remain small.

### 9 Additional experimental results

#### 9.1 Presomitic mesoderm aggregates (Exp. 2)

We repeated the analysis on a second presomitic mesoderm aggregate and obtained results consistent with those shown in the main text. The density pattern measure $P_\rho$ increases over time, and its temporal decomposition again shows that this increase is mainly driven by cell motility, with smaller contributions from cell division, boundary crossings, and cell disappearance (Fig. S8A–C).

The apparent expression pattern measure $P_{f\rho}$ also increases over time (Fig. S8D–F). However, the T/Brachyury expression maps do not show a clear large-scale expression pattern beyond the spatial structure already present in the cell density. Consistently, the decomposition of $P_{f\rho}$ is very similar to that of $P_\rho$, and the direct contribution of expression changes remains small. This suggests that, in this second aggregate, the apparent T/Brachyury expression pattern is mainly driven by cell rearrangements rather than by coordinated changes in T/Brachyury expression.

#### 9.2 Breast cancer cell lines (Exp. 2)

The second SUM159PT experiment confirms the division-dominated behavior observed in the main text, but with a weaker increase in pattern strength per cell. The density pattern measure $P_\rho$ increases over time, together with the emergence of spatial heterogeneity in cell density (Fig. S9A,B). Its decomposition shows that this increase is mainly driven by cell divisions, with an additional positive contribution from cell dis-/appearance, whereas motility and boundary crossings remain close to zero.

The subsampling analysis further indicates that the increase in $P_\rho$ is only weakly accompanied by an increase in the signal-to-noise ratio per cell, with $q_\rho$ changing only modestly between the first and last frames (Fig. S9C). Thus, in this experiment, the increase in $P_\rho$ is largely driven by the growth of the cell population, rather than by a strong amplification of density patterning per cell.

A similar behavior is observed for the apparent ALDH1A1 expression pattern. The measure $P_{f\rho}$ increases during the experiment, but the fitted values of $q_{f\rho}$ remain small (Fig. S9F). The decomposition shows that cell divisions provide the dominant positive contribution to $P_{f\rho}$, while changes in ALDH1A1 expression contribute negatively at late times (Fig. S9E). This supports the conclusion that the apparent ALDH1A1 pattern is mainly amplified through population growth and division-driven spatial structure, whereas expression dynamics do not reinforce the pattern.

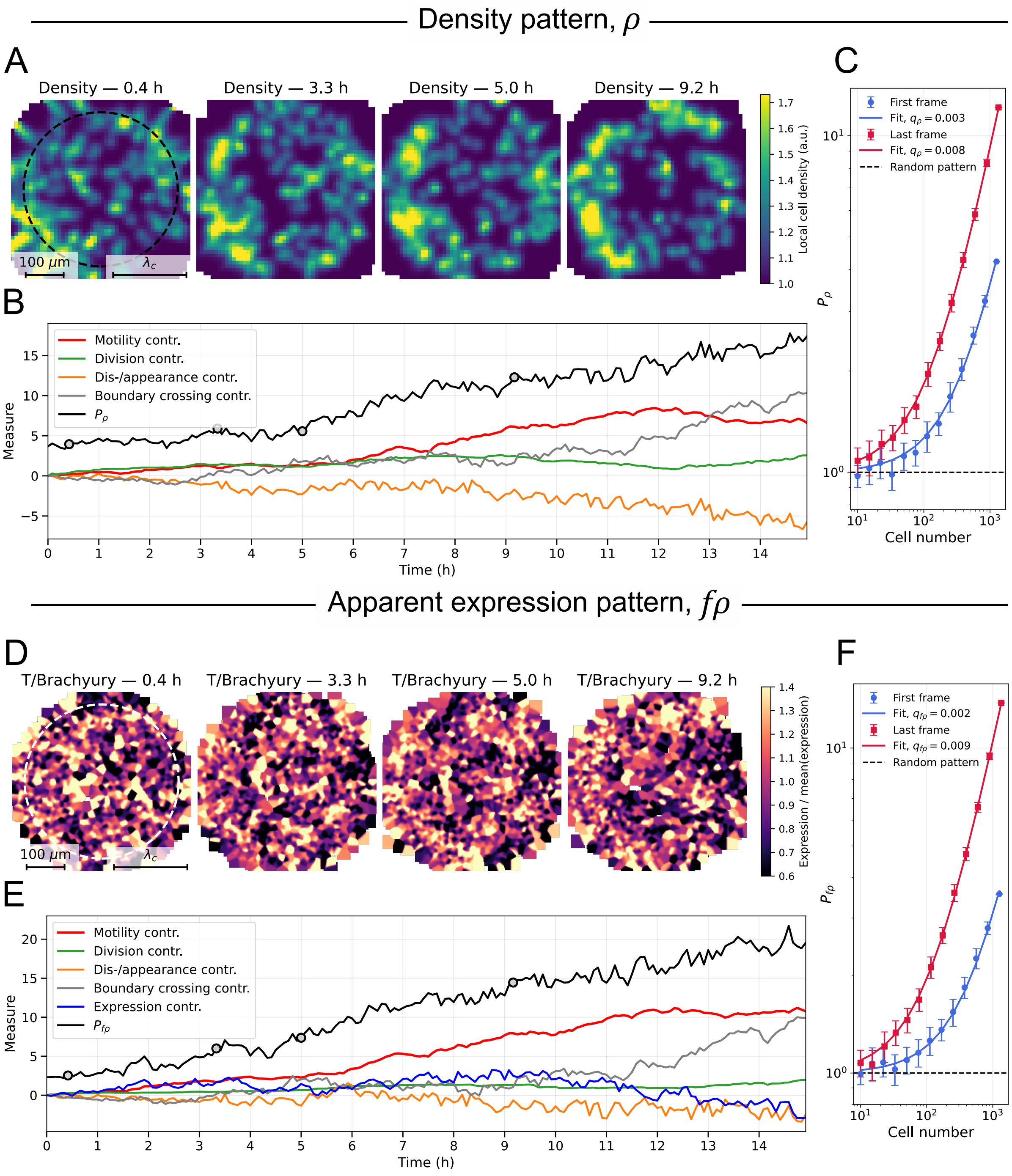

Figure S8: Pattern formation and cellular contributions in presomitic mesoderm cell aggregates (Exp. 2). (A) Spatial distribution of cell density in the same aggregate at four time points ($t = 0.4$, 3.3, 5.0 and 9.2 h). Density maps are reconstructed from tracked cell positions and shown as smoothed spatial fields for visualization. (B) Temporal evolution of the density pattern measure $P_\rho(t)$ together with the decomposition of $P_\rho(t) - P_\rho(0)$ into contributions by cell motility, cell division, boundary crossings, and cell disappearance. (C) Dependence of $P_\rho$ on population size $N$ obtained by subsampling, showing the temporal increase of the signal-to-noise ratio per cell. (D) Apparent T/Bra expression in the same aggregate at the same time points as in A, normalized by the spatial average. (E) Temporal evolution of the apparent expression pattern measure $P_{f\rho}(t)$ together with the decomposition of $P_{f\rho}(t) - P_{f\rho}(0)$ into contributions by cell motility, gene expression dynamics, cell division, boundary crossings, and cell disappearance. (F) Dependence of $P_{f\rho}$ on population size $N$, confirming the temporal increase of the signal-to-noise ratio per cell.

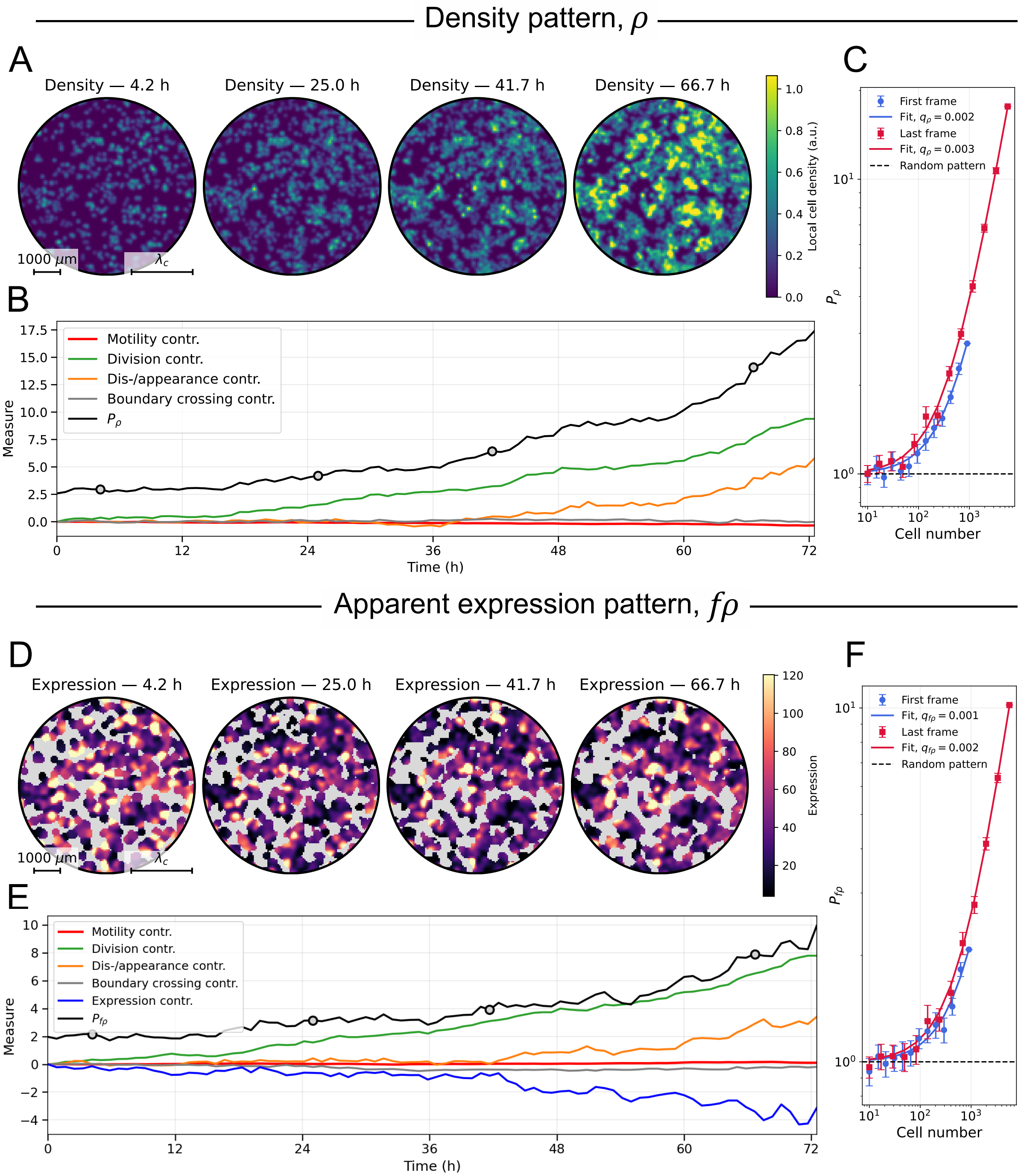

Figure S9: Pattern formation and cellular contributions in breast cancer cell populations (Exp. 2). (A) Spatial distribution of cell density at four time points ($t = 4.2$, 25.0, 41.7 and 66.7 h). Density maps are reconstructed from tracked cell positions and shown as smoothed spatial fields for visualization. (B) Temporal evolution of the density pattern measure $P_\rho(t)$ together with the decomposition of $P_\rho(t) - P_\rho(0)$ into contributions by cell motility, cell division, boundary crossings, and cell dis-/appearance. (C) Dependence of $P_\rho$ on population size $N$ obtained by subsampling, showing a weak increase of the signal-to-noise ratio per cell. (D) Apparent ALDH1A1 expression pattern at the same time points as in A, shown after normalization by the spatial average. (E) Temporal evolution of the apparent expression pattern measure $P_{f\rho}(t)$ together with the decomposition of $P_{f\rho}(t) - P_{f\rho}(0)$ into contributions by cell motility, gene expression dynamics, cell division, boundary crossings, and cell dis-/appearance. (F) Dependence of $P_{f\rho}$ on population size $N$, showing that the signal-to-noise ratio per cell remains small and changes only weakly during the observation window.

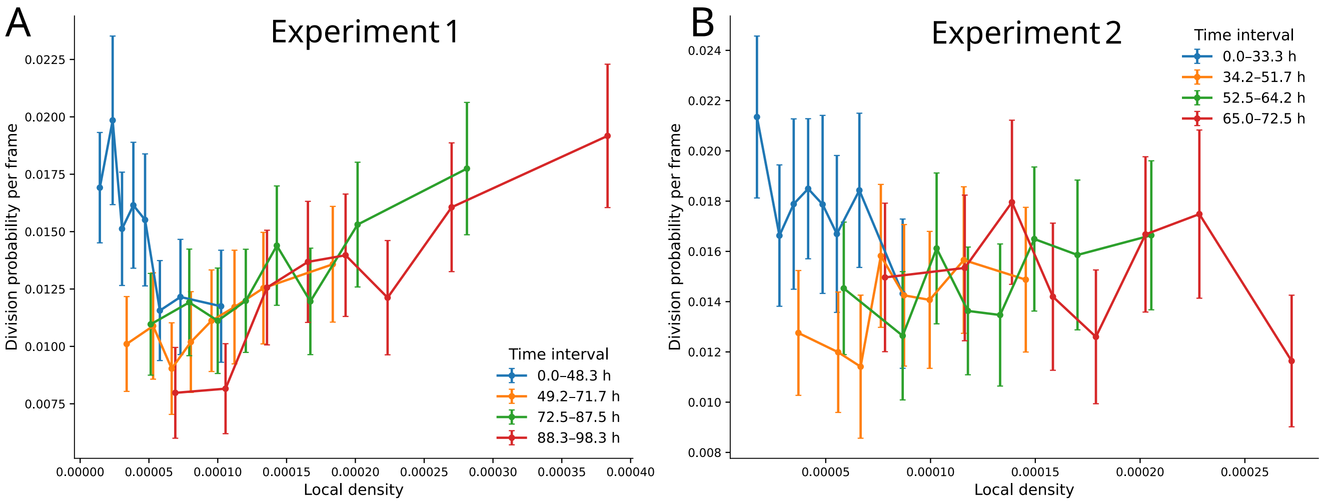

Figure S10: Density-dependent cell division in breast cancer cell populations. Local density was computed within a disk of radius $R = 338$ $\mu$m. Cells were divided into temporal groups containing equal numbers of cell-frame observations, ordered by time, and then binned by local density. (A) Division probability per frame as a function of local density in Experiment 1. (B) Same analysis for Experiment 2.

#### 10 Testing for density-dependent cell division in breast cancer cell populations

The increase in the signal-to-noise ratio per cell, $q_\rho$, in Experiment 1 suggests that cell divisions do not only contribute to increasing $P_\rho$ through an increase in cell number, but that they additionally sharpen the pattern.

To test whether cell divisions were biased toward dense regions, we computed for each cell the local density within a disk of radius $R = 338$ $\mu$m. Cells located less than the boundary of the field of view were excluded to avoid edge effects. We then recorded whether each cell divided during the following frame. Cells were divided into four temporal groups containing equal numbers of cell-frame observations, and the corresponding time intervals are shown in the legend. Within each temporal group, cells were binned according to local density, and the division probability per frame was computed in each bin.

In Experiment 1, division probability increased with local density at late times (Supplementary Fig. S10A), indicating that cells in denser regions were more likely to divide once the density pattern had emerged. This density-dependent bias is consistent with the positive contribution of cell division to the sharpening of the density pattern.

We applied the same analysis to Experiment 2 (Supplementary Fig. S10B). In this experiment, division probability did not show a clear monotonic increase with local density, including at late times. Thus, a clear density-dependent division bias was apparent in Experiment 1 but not in Experiment 2. This is consistent with the fact that $q_\rho$ remained very low in Experiment 2 (Fig. S9), suggesting that the density pattern in this experiment was mainly driven by the increase in cell number through divisions, without a strong bias favoring divisions in already dense regions.

#### 11 Code availability

To automatize computation, we developed the publicly available python tool CREDIT (Cellular Roles in Emergent Development Inferred from Tracks), which is available here.
